# RIOK2 phosphorylation by RSK promotes synthesis of the human small ribosomal subunit

**DOI:** 10.1101/2020.10.07.329334

**Authors:** Emilie L. Cerezo, Thibault Houles, Oriane Lié, Marie-Kerguelen Sarthou, Charlotte Audoynaud, Geneviève Lavoie, Maral Halladjian, Sylvain Cantaloube, Carine Froment, Odile Burlet-Schiltz, Yves Henry, Philippe P. Roux, Anthony K. Henras, Yves Romeo

## Abstract

Ribosome biogenesis lies at the nexus of various signaling pathways coordinating protein synthesis with cell growth and proliferation. This process is regulated by well-described transcriptional mechanisms, but a growing body of evidence indicates that other levels of regulation exist. Here we show that the Ras/mitogen-activated protein kinase (MAPK) pathway stimulates post-transcriptional stages of human ribosome synthesis. We identify RIOK2, a pre-40S particle assembly factor, as a new target of the MAPK-activated kinase RSK. RIOK2 phosphorylation by RSK promotes cytoplasmic maturation of late pre-40S particles, which is required for optimal protein synthesis and cell proliferation. RIOK2 phosphorylation facilitates its release from pre-40S particles and its nuclear re-import, prior to completion of small ribosomal subunits. Our results bring a detailed mechanistic link between the Ras/MAPK pathway and the maturation of human pre-40S particles, which open a hitherto poorly explored area of ribosome biogenesis.

## Introduction

Ribosomes are the universal macromolecular machines responsible for protein synthesis. Eukaryotic ribosomes consist of two subunits (40S and 60S) containing four ribosomal RNAs (rRNAs) and approximately 80 ribosomal proteins (RPs). Ribosome biogenesis begins in the nucleolus with the synthesis by RNA polymerase I (Pol I) of a polycistronic transcript precursor to the 18S, 5.8S and 28S rRNAs and of a precursor to the 5S rRNA by Pol III (Supplemental Fig. S1A). The nascent Pol I transcript is co-transcriptionally packaged into a pre-ribosomal particle that undergoes a series of maturation steps comprising folding and nucleolytic processing of the precursor RNA, chemical modifications of selected nucleotides, and incorporation of RPs and the 5S ribonucleoprotein particle (RNP). Early in the pathway, two discrete particles are generated, the pre-40S and pre-60S particles, precursors to the small and large subunits, respectively. These particles undergo independent maturation steps in the nucleolus and nucleoplasm, before being exported through the nuclear pore complexes. Once in the cytoplasm, pre-40S and pre-60S particles undergo final maturation steps before entering the pool of translation-competent subunits (Henras et al. 2015; Kressler et al. 2017).

Assembly and maturation of pre-ribosomes is promoted by scores of accessory factors associating transiently with the precursor particles and collectively referred to as assembly and maturation factors (AMFs) (Henras et al. 2015; Kressler et al. 2017; Aubert et al. 2018; Cerezo et al. 2019). Some of these factors participate in the structuring of pre-ribosomes via RNA-binding and/or protein-protein interaction domains, whereas others carry different enzymatic activities, such as nucleases, nucleotide modifying enzymes, putative RNA helicases, kinases, ATPases or GTPases. Ribosome AMFs have been rather exhaustively identified in yeast and the function of some of them is quite well characterized (Woolford and Baserga 2013). In human cells, ribosome synthesis appears more complex since large-scale studies revealed that it mobilizes an increased number of AMFs (Wild et al. 2010; Tafforeau et al. 2013; Badertscher et al. 2015; Farley-Barnes et al. 2018) and pre-rRNA processing involves a more complex series of events compared to yeast (Henras et al. 2015; Bohnsack and Bohnsack 2019).

Ribosome biogenesis is one of the most energetically demanding of cellular activities. This process needs to be tightly and dynamically regulated to accommodate cell growth and proliferation under favorable conditions, and to prevent energy waste under limiting conditions. The signal transduction cascades Ras/mitogen-activated protein kinase (MAPK) and phosphoinositide 3-kinase (PI3K)/AKT pathways activate ribosome biogenesis in response to various cues, such as growth factors, cellular energy status, and nutrient availability (Goodfellow and White 2007; Grummt 2003; Mahoney et al. 2009; Piazzi et al. 2019). Upon activation of the Ras/MAPK pathway (Supplemental Fig. S1B), the MAPKs ERK1/2 phosphorylate scores of substrates (Cargnello and Roux 2011), including members of the RSK (p90 ribosomal S6 kinase) family of protein kinases (Houles and Roux 2018; Romeo et al. 2012), which collectively promote cell growth and proliferation. The PI3K/AKT signaling pathway activates the mechanistic target of rapamycin (mTOR), which is a conserved Ser/Thr kinase that stimulates anabolic processes such as mRNA translation (Wullschleger et al. 2006; Roux and Topisirovic 2018). Sustained activation of these signaling molecules promotes ribosome production and increases global translation to feed the protein needs under conditions of growth and proliferation.

Activation of the MAPK and mTOR pathways promotes ribosome biogenesis by stimulating rDNA transcription (Kusnadi et al. 2015). ERK1/2 and RSK, as well as mTOR and its downstream target S6K (p70 ribosomal S6 kinase), phosphorylate major Pol I (RRN3/TIFI-A and UBF) and Pol III (TFIIIB) transcription factors, and thereby increase synthesis of the 47S pre-rRNA (Pol I) and the 5S rRNA (Pol III) (Zhao et al. 2003; Felton-Edkins et al. 2003; Sriskanthadevan-Pirahas et al. 2018; Stefanovsky et al. 2001, 2006a, 2006b; Stefanovsky and Moss 2008; Zhang et al. 2013). In addition, mTOR signaling stimulates transcription of genes encoding RPs and AMFs through the activity of S6Ks (Chauvin et al. 2014). Both pathways also promote translation of mRNAs encoding RPs and AMFs (Geyer et al. 1982; Asmal et al. 2003; Romeo et al. 2013).

By co-activating the syntheses of rRNAs, RPs and AMFs, MAPK and mTOR pathways participate in stoichiometric expression of ribosome components and ribosome synthesis machinery, and thereby coordinate the initial stages of ribosome production. Evidence in the literature suggests that these two pathways also regulate the post-transcriptional steps of ribosome synthesis. MAPK and mTOR pathways seem to promote pre-rRNA processing (Iadevaia et al. 2012; Isfort and Cody 1992), and mTOR inhibition induces the mislocalisation of several AMFs both in yeast and human cells (Vanrobays et al. 2008; Raman et al. 2014; Talkish et al. 2012; Honma et al. 2006). Although these data suggest that pre-ribosome assembly and maturation are also under the control of signaling pathways, no search has been undertaken so far to identify direct targets of the MAPK and mTOR signaling pathways among the large number of pre-ribosome AMFs.

Here we report that the MAPK pathway directly regulates discrete molecular events during the post-transcriptional steps of ribosome biogenesis, which fills a major gap between currently known functions of MAPK signaling in Pol I transcription and cytoplasmic translation. Our results unravel a link between RSK signaling and the maturation of human pre-40S particles, through the regulation of RIOK2, an atypical protein kinase of the RIO family involved in the synthesis of the small ribosomal subunit (LaRonde-LeBlanc and Wlodawer 2005).

## Results

### The MAPK pathway regulates post-transcriptional stages of ribosome biogenesis in a RSK-dependent manner

To assess whether the MAPK signaling pathway regulates pre-rRNA processing in human cells, we first examined the levels of various rRNA precursors to both the small and large ribosomal subunits (maturation pathway depicted in Supplemental Fig. S1A) upon pharmacological inhibition of ERK1/2 or RSK kinases. For this, serum-starved HEK293 cells were incubated with PMA (phorbol 12-myristate 13-acetate) to selectively stimulate the MAPK pathway prior to treatment with MEK1/2 (PD184352) or RSK (LJH685) inhibitors (Supplemental Fig. S1B). In this and all subsequent experiments, we have performed western blotting to assess the activation levels of ERK1/2 and RSK kinases (Fig. 1A, lower panels). Efficient activation of the MAPK pathway by PMA, serum or epidermal growth factor (EGF) was shown using phospho-specific antibodies targeted against phosphorylated ERK1/2 (T202/Y204) and RSK (S380). Using these tools, we found that treatment with MEK1/2 inhibitors (PD184352) abrogated ERK1/2 and RSK phosphorylation. As BI-D1870 and LJH685 target the N-terminal kinase domain of RSK, they do not prevent its phosphorylation at S380 by the C-terminal kinase domain. Treatments with both MEK1/2 and RSK inhibitors induced changes in the processing pathway leading to the production of the 18S rRNA, the RNA component of the small ribosomal subunit. Inhibition of MEK1/2 induced a strong accumulation of the 30S precursor and a marked reduction in the production of all downstream intermediates, in particular the 18S-E pre-rRNA, which is the ultimate precursor to the mature 18S rRNA (Fig. 1A and 1B). Inhibition of RSK resulted in a milder accumulation of the 30S precursor, without affecting the accumulation of the 21S and 18S-E precursors. With regards to the large ribosomal subunit processing pathway, accumulation of the 32S precursor was not strongly affected by MAPK pathway inhibition, and inhibition of ERK1/2, but not RSK, resulted in a decreased accumulation of the 12S precursor (Supplemental Fig. S1C and S1D).

**Figure 1.**
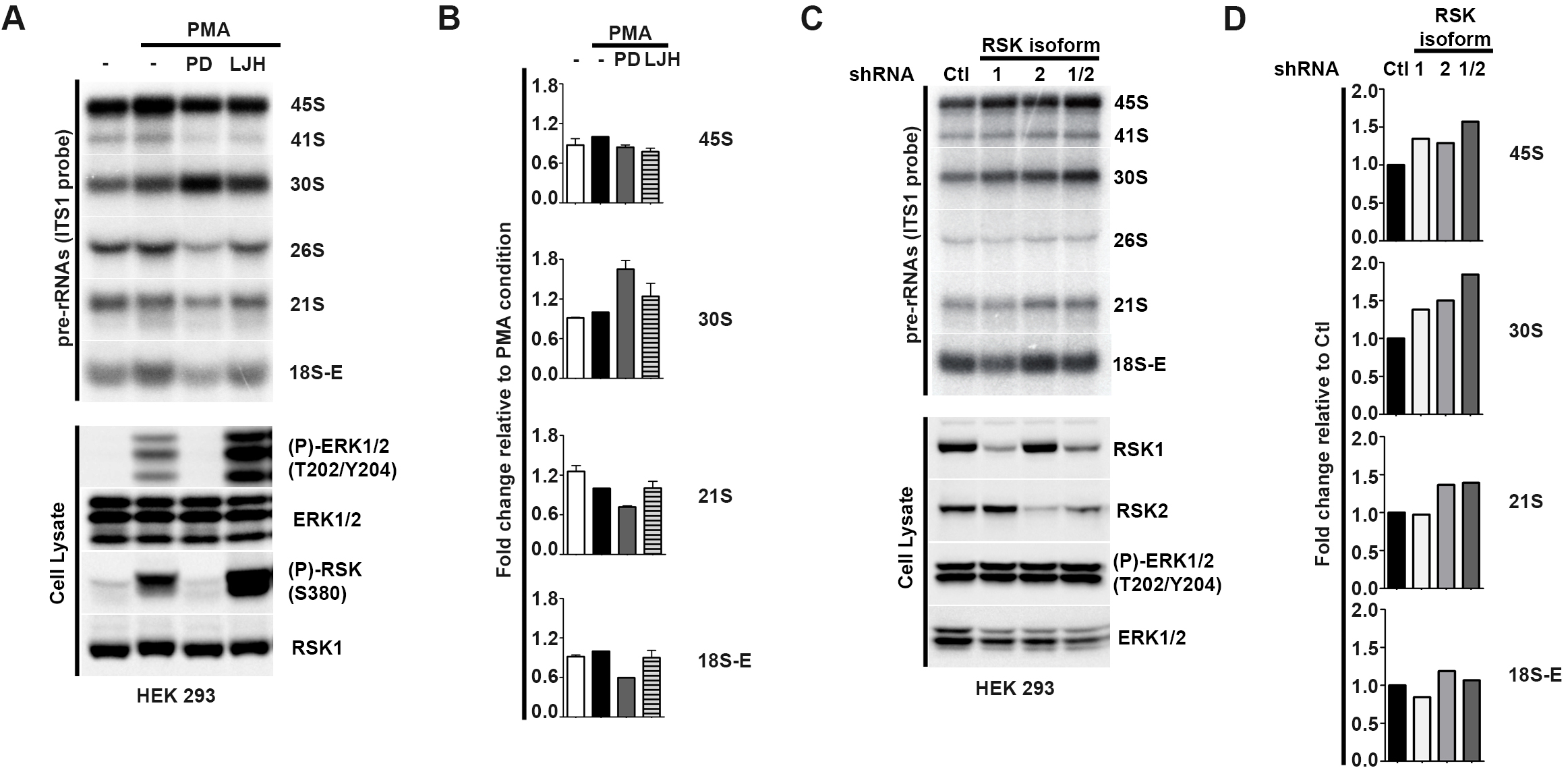
The MAPK signaling pathway regulates post-transcriptional stages of ribosome biogenesis in a RSK-dependent manner. *(A)* Serum-starved HEK293 cells were stimulated with PMA (100 ng/ml) for 30 min then treated with 10 µM of PD184352 or LJH685 inhibitors for 3h. Total RNAs were extracted and pre-rRNA levels were monitored by Northern blot (NB, see Supplemental Fig. S1 for a detailed representation of the human pre-rRNA processing pathway and position of the probes). Total proteins were extracted and analyzed by Western blot (WB) using indicated antibodies. *(B)* Quantification of pre-rRNA levels obtained in *(A)* using MultiGauge software (Fujifilm). Graphic representations were expressed as fold changes. *(C)* pre-rRNA and protein levels were monitored as in *(A)* from HEK293 cells expressing shRNA targeting an irrelevant sequence (Ctl) or RSK1 (1), RSK2 (2) or RSK1 and RSK2 (1/2). *(D)* Quantification of pre-rRNA levels as in *(C)*.

The RSK family comprises four closely related Ser/Thr kinases (RSK1-4) expressed from independent genes. Both RSK1 and RSK2 promote cell growth and proliferation, and are the predominant RSK isoforms expressed in HEK293 cells. To validate the implication of RSK in pre-ribosome maturation, we generated HEK293 stable cell lines expressing shRNAs specifically targeting RSK1 and/or RSK2 (Fig. 1C and 1D). We confirmed RSK1/2 depletion efficiency by western blot (Fig. 1C, lower panels) and further detected high levels of ERK phosphorylation upon RSK knockdown, indicating that the MAPK pathway upstream of RSK is still active. Like LJH685 treatment, knockdown of RSK1, RSK2 or both RSK1 and RSK2 increased the levels of the 30S precursor, with more variable effects on the 21S and 18S-E pre-rRNAs, which could reflect a difference in knockdown efficiency and/or slightly different functions of the isoforms in pre-rRNA processing (Fig. 1C and 1D). RSK knockdown but not LJH685 treatment resulted in the accumulation of the 45S precursor, but we suspect that this discrepancy rather reflects a difference in the growth conditions, namely acute response (PMA treatment of serum-starved cells) for LJH685 treatment versus steady state growth in serum-containing medium for knockdown experiments. As with pharmacological inhibitors, knockdown of RSK1, RSK2 or both isoforms did not induce major changes in the accumulation of the large subunit intermediates 32S and 12S (Supplemental Fig. S1E and S1F).

Altogether, these data strongly suggest that the MAPK pathway influences the kinetics of some stages of pre-ribosome assembly and maturation in human cells, and that RSK participates in this process, possibly by modulating the activity of selected pre-ribosome AMFs.

### RIOK2 is phosphorylated at Ser483 upon activation of the MAPK pathway

To understand the precise mechanisms by which RSK regulates post-transcriptional stages of ribosome synthesis, we performed an *in silico* screen to identify potential RSK phosphorylation targets involved in this process. Using the Scansite bioinformatics tool (https://scansite4.mit.edu/4.0/), we searched for the canonical Arg/Lys-X-Arg/Lys-X-X-pSer/Thr (RXRXXpS/T) motif found in RSK substrates, within the sequences of an exhaustive list of human AMFs potentially involved in the synthesis of the 40S subunit (Badertscher et al. 2015; Tafforeau et al. 2013; Woolford and Baserga 2013). We found a number of AMFs featuring high confidence RXRXXpS/T phosphorylation motifs, among which the RIOK2 protein kinase. RIOK2 is a 552 amino acid protein containing two RXRXXpS/T sites in the C-terminal region of the protein, a high-stringency site predicting phosphorylation at Ser483 (S483) and a medium-stringency site at Thr481 (T481) (Supplemental Fig. S2A).

To determine whether RIOK2 is phosphorylated at RXRXXpS/T motifs and whether this phosphorylation event responds to MAPK pathway activation, HEK293 cells transiently expressing HA-tagged RIOK2 were serum-starved and stimulated with EGF or PMA agonists. Phosphorylation of immunoprecipitated HA-RIOK2 was analyzed by immunoblotting using antibodies detecting the phosphorylated consensus RXRXXpS/T motif (Fig. 2A). Notably, we found that activation of the MAPK pathway clearly stimulated RIOK2 phosphorylation at the RXRXXpS/T motif. Pre-treatment of starved cells with MEK1/2 inhibitor (PD184352) abrogated the induction of RIOK2 phosphorylation in response to EGF or PMA, indicating that RIOK2 is phosphorylated on RXRXXpS/T consensus sites in a MAPK-dependent manner.

**Figure 2.**
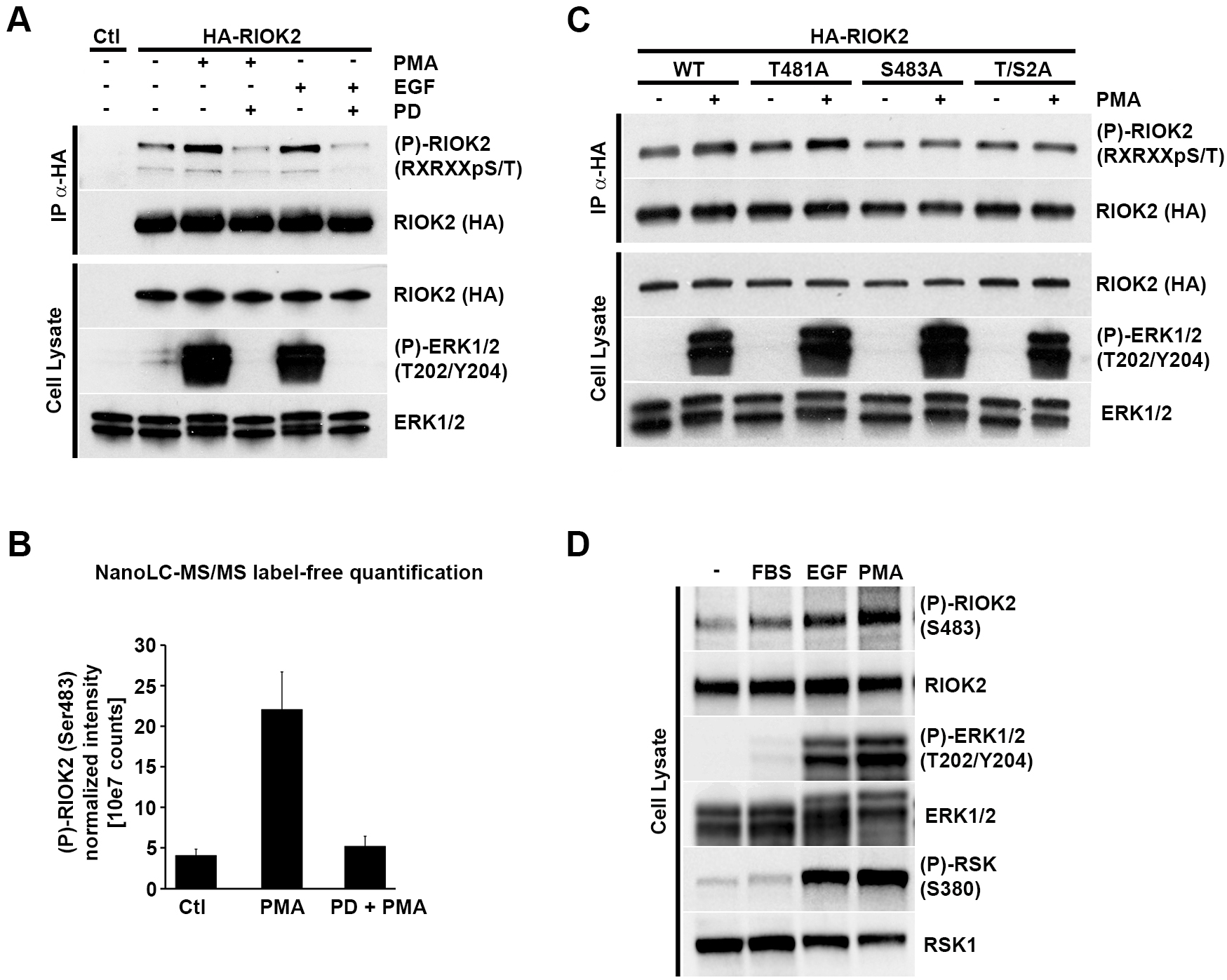
RIOK2 is phosphorylated at Ser483 upon activation of the MAPK pathway. *(A)* HEK293 cells were transfected with a plasmid expressing HA-RIOK2 or an empty vector (Ctl). HA-RIOK2 was immunoprecipitated from serum-starved cells treated or not with PD184352 for 1h (10 µM) prior to PMA (100 ng/ml, 20 min) or EGF (25 µg/ml, 10 min) stimulation. Samples were analyzed by WB using anti-RXRXXpS/T or anti-HA antibodies. *(B)* HA-RIOK2 was immunoprecipitated from serum-starved HEK293 cells treated or not with PD184352 (10 µM, 1h) prior to PMA stimulation (100 ng/ml, 20 min). Purified HA-RIOK2 was isolated following SDS-PAGE, in gel digested with trypsin and the resulting peptides were submitted to nano-LC-MS/MS analysis. Label-free quantitative analysis of phosphorylation of the different RIOK2 phospho-peptides was performed as specified in *Method Details*. Data are representative of triple biological replicate experiments for each condition. *(C)* HEK293 cells expressing HA-tagged versions of WT or mutant versions of RIOK2 T481A, S483A or S/T2A were serum starved prior to PMA stimulation (100 ng/ml, 20 min). HA-RIOK2 was immunoprecipitated and samples were analyzed by WB as in *(A). (D)* Serum-starved HEK293 cells were stimulated with different agonists of the MAPK pathway. Phosphorylation of endogenous RIOK2 was monitored by WB using antibodies against phosphorylated RIOK2 Ser483.

We next attempted to identify RIOK2 phosphorylation sites that are regulated by MAPK signaling using a label-free quantitative mass spectrometry (MS) approach. HEK293 cells expressing HA-tagged RIOK2 were serum-starved overnight and pre-treated or not with MEK1/2 inhibitor (PD184352) prior to PMA stimulation (Supplemental Fig. S2B). Immunoprecipitated HA-RIOK2 was isolated using SDS-PAGE and digested in-gel with trypsin. Samples were analyzed by nano-liquid chromatography-tandem MS (nanoLC-MS/MS) and database searched for putative post-translational modifications, such as phosphorylation. The relative abundance of all identified phosphopeptides corresponding to 12 phosphorylation sites was evaluated (Table S1), amongst which Ser483 was the only RIOK2 residue whose phosphorylation robustly increased upon PMA stimulation and returned to basal levels upon MEK1/2 inhibition (Fig. 2B and Supplemental Fig. 2C). Phosphorylation at Thr481 was not detected, suggesting that this residue is not phosphorylated in HEK293 cells. Interestingly, the RSK phosphorylation motif containing Ser483 is conserved within vertebrates, suggesting that it is involved in an important biological function (Supplemental Fig. S2D).

To further confirm that the MAPK pathway induces RIOK2 phosphorylation at Ser483, we transiently expressed in HEK293 cells HA-tagged versions of WT or mutant RIOK2 in which Ser483 and/or Thr481 were substituted for a non-phosphorylatable alanine (RIOK2^S483A^, RIOK2^T481A^, RIOK2^T481A/S483A^ (Fig. 2C). These cells were serum-starved, stimulated with PMA and HA-RIOK2 was purified from cell lysates and analyzed by immunoblotting. Mutation of Ser483 alone or in combination with Thr481 abolished RIOK2 phosphorylation after PMA stimulation. Since mutation of Thr481 alone did not reduce RIOK2 phosphorylation and since we did not detect Thr481 phosphorylation *in vivo*, we conclude that activation of MAPK signaling induces RIOK2 phosphorylation specifically at Ser483.

To validate these results, we generated antibodies specifically directed against the Ser483-phosphorylated RIOK2 peptide and monitored the phosphorylation status of endogenous RIOK2 in serum-starved HEK293 cells or in response to agonists of the MAPK pathway (Fig. 2D). Phosphorylation of RIOK2 at Ser483 clearly increased upon stimulation with serum, EGF or PMA, demonstrating that activation of MAPK signaling results in phosphorylation of endogenous RIOK2 at Ser483.

### RIOK2 is a direct RSK substrate

To test whether RSK is responsible for RIOK2 phosphorylation at Ser483, we treated serum-starved HEK293 cells with PMA, with or without prior treatment with the MEK1/2 (PD184352) or RSK (BI-D1870 and LJH685) inhibitors and analyzed RIOK2 phosphorylation at S483 (Fig. 3A). Consistent with a role for RSK in RIOK2 phosphorylation, we observed that treatment of cells with each of these inhibitors abrogated Ser483 phosphorylation upon PMA stimulation.

**Figure 3.**
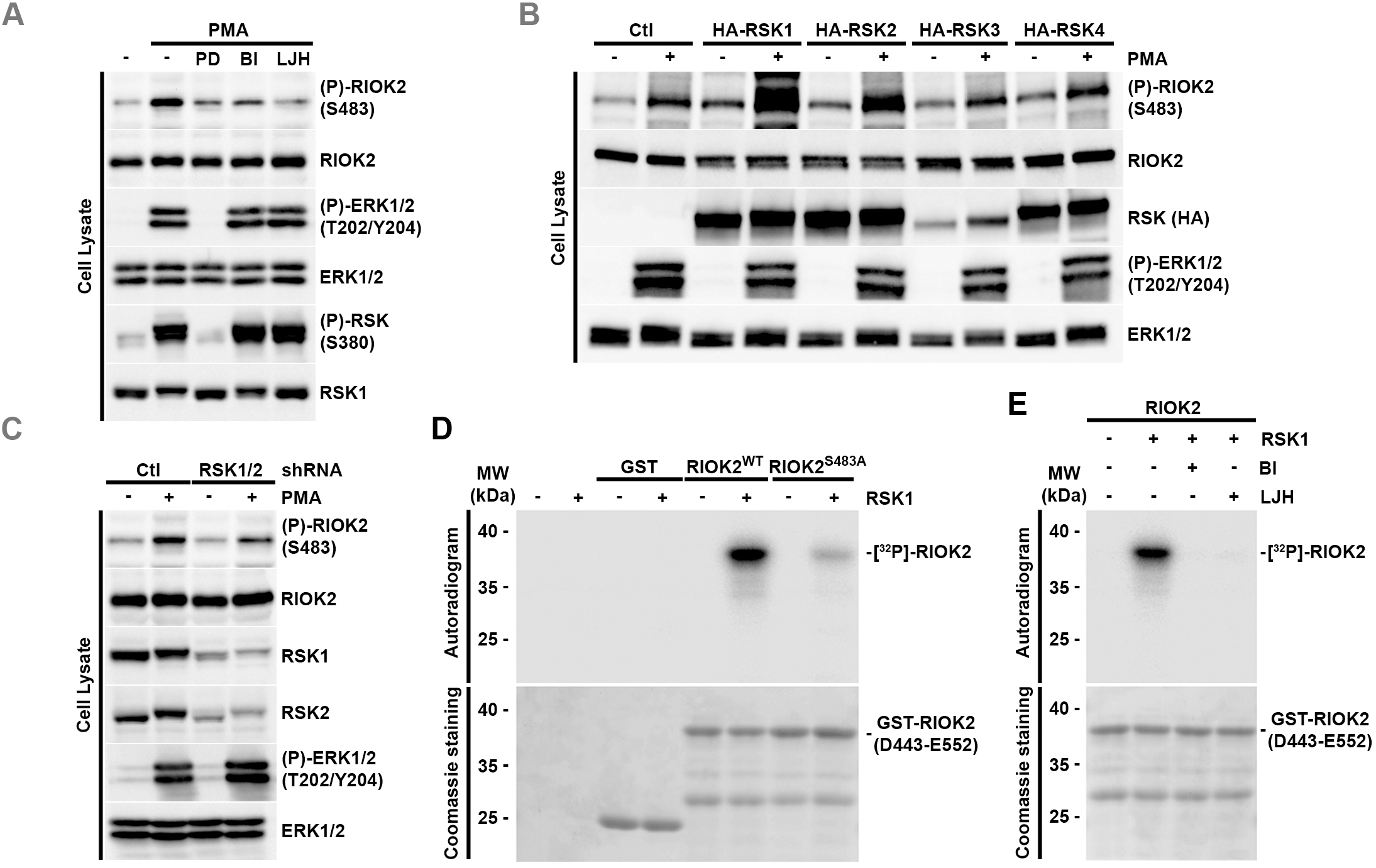
RIOK2 is a direct RSK substrate. *(A)* Serum-starved HEK293 cells were treated with PD184352, BI-D1870 or LJH685 (10 µM) for 1 h prior to PMA stimulation (100 ng/ml, 20 min). Phosphorylation of endogenous RIOK2 at Ser483 was monitored by WB using specific antibodies. *(B)* HEK293 cells were transfected with vectors over-expressing HA-tagged RSK1, RSK2, RSK3 or RSK4, or the empty vector (Ctl). Following serum-starvation and PMA stimulation (100 ng/ml, 20 min), samples were analyzed by WB using indicated antibodies. Note: RSK3 is overexpressed at the same level as the other RSK isoforms but, as previously reported (Zhao et al. 1996), the protein is largely insoluble and is pelleted during the centrifugation step to yield the total lysate. *(C)* HEK293 cells expressing shRNA targeting an irrelevant sequence (Ctl) or both RSK1 and RSK2 (RSK1/2) were processed as in *(B). (D)* Activated RSK1 was incubated in the presence of γ[^32^P]-ATP with either GST alone, or a GST-RIOK2 peptide (D443-E552) containing either S483 (RIOK2^WT^) or the non-phosphorylatable version (RIOK2^S483A^). The resulting samples were analyzed by SDS-PAGE and revealed by autoradiography or Coomassie blue staining. *(E)* As in *(D)* in the presence or not of RSK inhibitors BI-D1870 or LJH685 (10 mM).

As four RSK isoforms are expressed in human cells, we next investigated which one(s) is/are involved in RIOK2 phosphorylation at Ser483. HEK293 cells over-expressing each of the four RSK isoforms were stimulated with PMA and analyzed for RIOK2 phosphorylation at Ser483 by immunoblotting (Fig. 3B). Our results show that overexpression of RSK1 or RSK2, but not RSK3 or RSK4, increased RIOK2 phosphorylation at Ser483 upon PMA treatment, suggesting that the former are the predominant RSK isoforms involved in the regulation of RIOK2. The role of RSK1 and RSK2 was further confirmed using stable (shRNAs) or transient (siRNAs) RNA interference. Consistent with a dual role for RSK1 and RSK2, we found that knockdown of both isoforms resulted in a clear reduction of RIOK2 Ser483 phosphorylation upon activation of MAPK signaling, compared to control cells (Fig. 3C and Supplemental Fig. S3, respectively).

To determine if RIOK2 is a direct RSK substrate, we performed *in vitro* phosphorylation assays using a GST-tagged C-terminal fragment of RIOK2 spanning residues Asp443 to Glu552 purified from *E. coli* (Fig. 3D and 3E). Upon incubation with active RSK1 purified from Sf9 insect cells and γ[^32^P]-ATP, this peptide became efficiently phosphorylated (Fig. 3D). As specificity controls, we showed that GST alone or a mutant peptide in which Ser483 was replaced by a non-phosphorylatable alanine were poorly phosphorylated by RSK. Addition of ATP-competitive RSK inhibitors (BI-D1870 or LJH685) to the *in vitro* assay fully compromised [^32^P] incorporation into the RIOK2 peptide, demonstrating that this event requires RSK catalytic activity (Fig. 3E). Together, these experiments strongly suggest that RSK directly promotes RIOK2 phosphorylation at Ser483.

### RIOK2 phosphorylation at Ser483 is required for efficient maturation of pre-40S particles

Both yeast Rio2 and human RIOK2 are essential for cell viability (Vanrobays et al. 2003; Asquith et al. 2019). In agreement with this, RIOK2 is required for cell proliferation, migration and survival of glioblastoma (Read et al. 2013; Song et al. 2020). RIOK2 has been suggested to function in mitotic progression (Liu et al. 2011) but its best documented molecular function is linked to the synthesis of the small ribosomal subunit. Depletion of Rio2/RIOK2 prevents processing of the last precursor to the mature 18S rRNA (20S pre-rRNA in yeast or 18S-E pre-rRNA in human) within cytoplasmic pre-40S particles and therefore inhibits production of the 40S subunit (Zemp et al. 2009; Rouquette et al. 2005; Vanrobays et al. 2003). The precise function of RIOK2 in the maturation of pre-40S particles remains ill-defined. Yeast Rio2 features autophosphorylation and ATPase activities *in vitro* and has been suggested to function as an ATPase in the maturation process rather than a *bona fide* kinase (Ferreira-Cerca et al. 2012). In contrast, human RIOK2 was recently shown to phosphorylate DIM1 *in vitro*, which is a component of nuclear pre-40S particles (Sloan et al. 2019). Human RIOK2 forms catalytically inactive homodimers *in vitro*, suggesting that some aspects of RIOK2 regulation *in vivo* may involve dimerization (Maurice et al. 2019).

To assess the functional relevance of RIOK2 phosphorylation at Ser483, we used a CRISPR/Cas9 knock-in approach to generate human eHAP1 haploid cell lines expressing mutant versions of RIOK2, bearing a substitution of Ser483 for either a non-phosphorylatable alanine (RIOK2^S483A^) or a phosphomimetic aspartic acid (RIOK2^S483D^) (Agudelo et al. 2017). Notably, we found that RIOK2^S483A^ mutant cell lines displayed a significantly decreased proliferation rate, as assessed by both MTS assay (Fig. 4A) and cell counting (Supplemental Fig. S4A), indicating that phosphorylation of RIOK2 at Ser483 is required for optimal cell proliferation. This proliferation defect was found to be less pronounced than that resulting from treatment with PD184352 or BI-D1870 inhibitors (Supplemental Fig. S4B), likely because the latter affect several MAPK-dependent cellular processes. We further observed that the slower proliferation rate of RIOK2^S483A^-expressing mutant cells is not correlated with a significant increase in cell death mechanisms, such as apoptosis (Supplemental Fig. S4C). Interestingly, abrogation of RIOK2 phosphorylation at Ser483 impairs global protein synthesis, as measured using the surface sensing of translation (SUnSET) method (Schmidt et al. 2009) (Fig. 4B and Supplemental Fig. S4D). Since RIOK2 functions in the last stages of pre-40S particle maturation, these results suggest that RIOK2 phosphorylation at Ser483 is important for production of translation-competent ribosomes.

**Figure 4.**
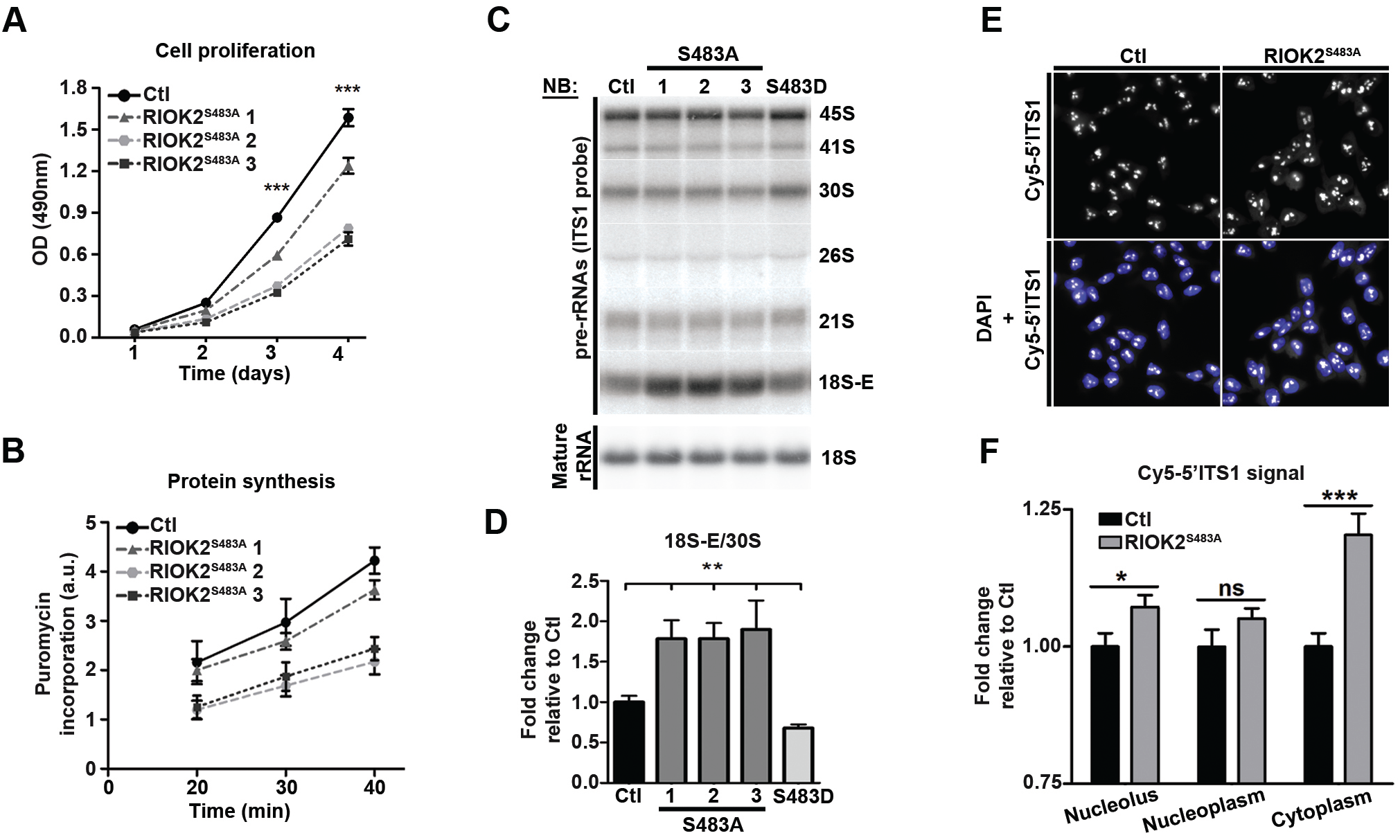
RIOK2 phosphorylation at Ser483 is required for efficient maturation of pre-40S particles. *(A)* MTS assays were performed on Control (Ctl) and RIOK2^S483A^ (1 to 3) eHAP1 cell lines at the indicated time points. ODs at 490 nm were measured using Spectramax. *(B)* Control (Ctl) and RIOK2^S483A^ (1 to 3) eHAP1 cell lines were incubated with 1 µM puromycin for the indicated times. Levels of puromycin-labelled peptides were monitored by WB using anti-puromycin antibodies. WB signals were quantified using ImageLab software and expressed as arbitrary units (a.u.). *(C), (D)*, Total cellular RNAs were extracted from control (Ctl), RIOK2^S483A^ (1 to 3) or RIOK2^S483D^ eHAP1 cell lines. Accumulation levels of pre-rRNAs and mature rRNAs were analyzed by NB and quantified as in Fig. 1, n=3. *(E)* FISH experiments performed on RIOK2^WT^ (Ctl) and RIOK2^S483A^ eHAP1 cell lines. Pre-rRNAs were detected using a Cy5-labeled 5’-ITS1 probe. Cells were stained with DAPI to visualize nuclei, and images were captured in identical conditions. *(F)* Nucleolar, nuclear and cytoplasmic fluorescence signals were quantified using ImageJ software. Graph representations show fold changes in RIOK2^S483A^ relative to RIOK2^WT^ cell line (n=100 cells from different fields). Statistically significant differences are indicated by asterisks (***: P≤0.001, **: P≤0.01, *: P≤0.05, One-tailed Mann Whitney test).

To delineate the molecular mechanism at the origin of this phenotype, we analyzed pre-rRNA processing in RIOK2^WT^, RIOK2^S483A^ and RIOK2^S483D^ cell lines. Total RNAs were extracted from these cells and levels of rRNA precursors were analyzed by Northern blotting (Fig. 4C upper panel). Interestingly, we found a ∼2-fold accumulation of the 18S-E precursor in all cell lines expressing RIOK2^S483A^ (Fig. 4C and 4D), indicating that the maturation of pre-40S particles is affected by the loss of Ser483 phosphorylation. RIOK2 knockdown in eHAP1 cells resulted in a similar accumulation of 18S-E pre-rRNA (Supplemental Fig. S4E and S4F), suggesting that Ser483 plays an important role in the regulation of RIOK2 function. In contrast, accumulation of the 18S-E pre-rRNA was not remarkably affected in cells expressing RIOK2^S483D^ despite a slight increase in the accumulation of the 45S and 30S precursors (Fig. 4C and 4D). No change in mature 18S rRNA levels was observed in these cell lines (Fig. 4, Supplemental Fig. S4G and S4H), suggesting that the steady state levels of mature 40S ribosomal subunits is not altered.

Pre-40S particles containing the 18S-E pre-rRNA are generated in the nucleolus (Nieto et al. 2020). They undergo maturation steps in the nucleoplasm before being exported to the cytoplasm, where final maturation events lead to the production of the mature 18S rRNA of the 40S subunit (Cerezo et al. 2019). To delineate more precisely which stage of pre-40S particle maturation is delayed in RIOK2^S483A^ cells, we performed fluorescence *in situ* hybridization (FISH) experiments using a probe detecting all pre-rRNAs of the small subunit maturation pathway (ITS1 probe, Supplemental Fig. 1A), including the 18S-E pre-rRNA (Fig. 4E and 4F). Cells expressing RIOK2^S483A^ displayed a strong accumulation of the 18S-E precursor in the cytoplasm compared to control cells, indicating that a significant proportion of pre-40S particles whose maturation is delayed accumulates in the cytoplasm. Collectively, our results strongly suggest that phosphorylation of RIOK2 at Ser483 is important for late, cytoplasmic stages of pre-40S particle maturation.

### Loss of RIOK2 phosphorylation at Ser483 increases its association with cytoplasmic pre-40S particles

RIOK2 is incorporated into pre-40S particles in the nucleus, participates to their export through direct binding to the CRM1 exportin, and dissociates from cytoplasmic pre-40S particles to get recycled back into the nucleus (Zemp et al. 2009; Fischer et al. 2015; Vanrobays et al. 2003). Rio2/RIOK2 catalytic activity contributes to its recycling into the nucleus (Geerlings et al. 2003; Zemp et al. 2009; Ferreira-Cerca et al. 2012; Knüppel et al. 2018). Furthermore, a defect in RIOK2 release is correlated with aberrant retention within pre-40S particles of other late AMFs, such as ENP1/Bystin, PNO1/DIM2, LTV1 and NOB1, the endonuclease responsible for conversion of the 18S-E precursor into mature 18S rRNA (Zemp et al. 2009).

To elucidate the molecular impact of RSK-dependent RIOK2 phosphorylation during the maturation of pre-40S particles, we first compared the nucleo-cytoplasmic distribution of RIOK2^WT^ and RIOK2^S483A^ using immunofluorescence (IF) microscopy. Our results indicate that RIOK2^S483A^ accumulates in the cytoplasm to a greater extent than RIOK2^WT^ (Fig. 5A). Quantification of the nuclear and cytoplasmic signals revealed a significant increase in the cytoplasmic to nuclear localization ratio of RIOK2^S483A^ compared to RIOK2^WT^ (Fig. 5B). These data indicate that in RIOK2^S483A^-expressing cells, both the mutant RIOK2 protein and pre-40S particles (Fig. 4E and 4F) accumulate in the cytoplasm.

**Figure 5.**
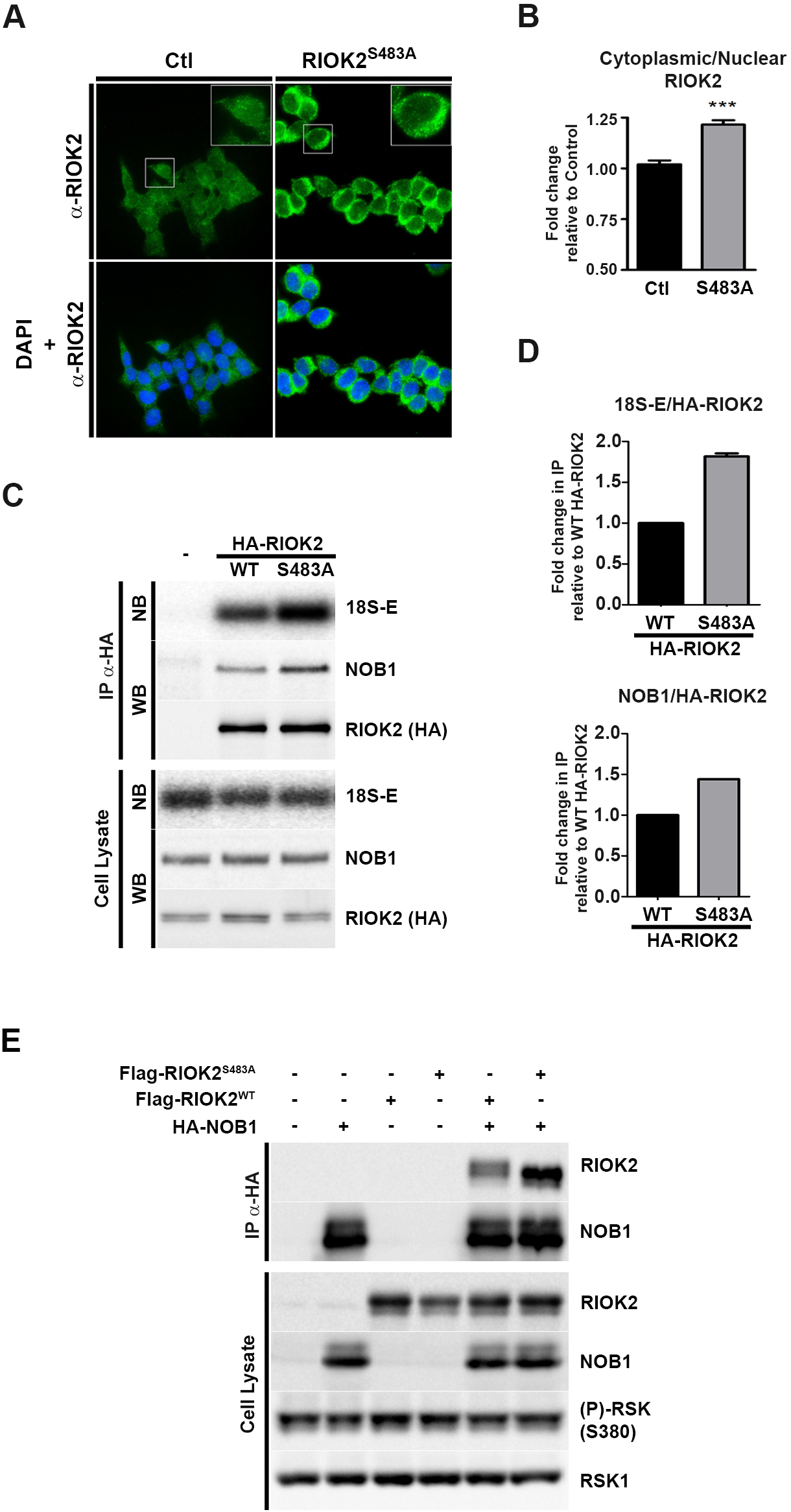
Loss of RIOK2 phosphorylation at Ser483 increases its association with cytoplasmic pre-40S particles. *(A)* RIOK2 localization was analyzed by IF microscopy using anti-RIOK2 antibodies in RIOK2^WT^ (Ctl) and RIOK2^S483A^ eHAP1 cell lines. Nuclei were visualized by DAPI staining. *(B)* Quantification of fluorescence observed in *(A)* using ImageJ software expressed as fold-change (n=100 cells from different fields). Statistically significant differences are indicated by asterisks (***: P<0.0001, One-tailed Mann Whitney test). *(C)* HEK293 cells were transfected with plasmids expressing HA-tagged RIOK2^WT^ or RIOK2^S483A^, or with empty vector (-). HA-RIOK2 was immunoprecipitated and co-immunoprecipitated proteins and 18S-E pre-rRNA were analyzed by WB and NB, respectively. *(D)* Quantification of the WB and NB signals obtained in IPs from *(C)* expressed as fold-changes compared to immunoprecipitated HA-RIOK2^WT^. *(E)* HEK293 cells were co-transfected with plasmids expressing HA-NOB1 and either Flag-RIOK2^WT^ or Flag-RIOK2^S483A^. HA-NOB1 was immunoprecipitated and the co-immunoprecipitated proteins were analyzed by WB.

To more directly assess the physical association between RIOK2 and pre-40S particles, we performed immunoprecipitation (IP) experiments. We purified pre-40S particles from HEK293 cells expressing HA-tagged versions of RIOK2^S483A^ or RIOK2^WT^, and quantified the levels of co-purified 18S-E pre-rRNA by Northern blotting. Using this approach, we found that the 18S-E pre-rRNA co-immunoprecipitated with ∼2-fold increased efficiency with RIOK2^S483A^ compared to RIOK2^WT^ (Fig. 5C and 5D). Importantly, ectopic expression of RIOK2^S483A^ did not change the global level of 18S-E pre-rRNA (Fig. 5C, Cell Lysate), most likely due to the presence of endogenous RIOK2. Endogenous NOB1 also co-immunoprecipitated with increased efficiency with RIOK2^S483A^ (Fig. 5C and 5D). These results were confirmed by immunoprecipitating HA-tagged NOB1 from HEK293 cells also expressing Flag-tagged versions of either RIOK2^WT^ or RIOK2^S483A^ (Fig. 5E). We observed that Flag-RIOK2^S483A^ was more efficiently co-purified with HA-NOB1 compared to Flag-RIOK2^WT^. We conclude that the loss of RSK-mediated phosphorylation at Ser483 increases the steady-state association of RIOK2 with pre-40S particles.

### RIOK2 phosphorylation at Ser483 facilitates its release from pre-40S particles and re-import into the nucleus

To understand the molecular determinants accounting for the higher steady-state association of RIOK2^S483A^ with pre-40S particles, we performed *in vitro* RIOK2 dissociation assays from purified pre-40S particles (Fig. 6A). For this, we purified pre-40S particles using HA-NOB1 from HEK293 cells also expressing Flag-tagged versions of either RIOK2^WT^, RIOK2^S483A^ or RIOK2^S483D^. Pre-40S particles bound to the anti-HA affinity matrix were then incubated in a buffer intended to promote RIOK2 catalytic activity, which is required for its dissociation from pre-40S particles (Zemp et al. 2009; Knüppel et al. 2018). Using this approach, we found that the dissociation kinetics of RIOK2^S483A^ was significantly less efficient than RIOK2^WT^, as only ∼30% of the mutant protein was dissociated from pre-40S particles compared to RIOK2^WT^ following a 90-min incubation (Fig. 6B). In contrast, the phosphomimetic RIOK2^S483D^ mutant was found to display dissociation kinetics quite similar to RIOK2^WT^, since by 90 min of incubation, the same amounts of RIOK2^S483D^ and RIOK2^WT^ were released from pre-40S particles. We concluded from these experiments that RIOK2^S483A^ is more stably associated to pre-40S particles compared to RIOK2^WT^ or RIOK2^S483D^, which strongly suggests that phosphorylation of RIOK2 at Ser483 facilitates its release from cytoplasmic pre-40S particles.

**Figure 6.**
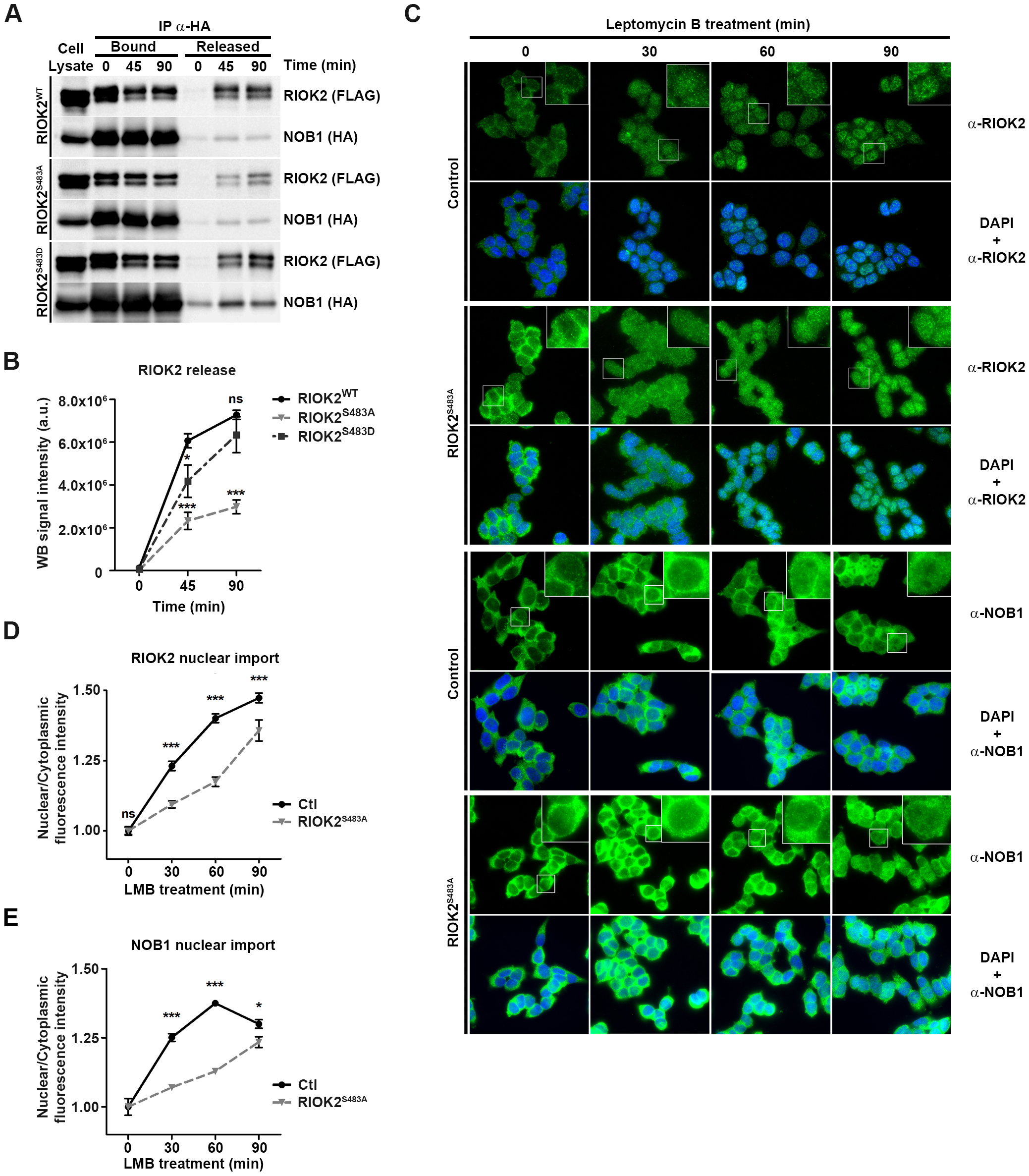
RIOK2 phosphorylation at Ser483 facilitates its release from pre-40S particles and re-import into the nucleus. *(A)* HEK293 cells were co-transfected with plasmids expressing HA-NOB1 and Flag-RIOK2^WT^, Flag-RIOK2^S483A^ or Flag-RIOK2^S483D^. Pre-40S particles were immunopurified via HA-NOB1 and subsequently incubated for 45 or 90 min with a buffer inducing RIOK2 release at 16°C. RIOK2 proteins in supernatant (released proteins) and retained on beads (pre-40S-bound) were analyzed by WB. Experiments with Flag-RIOK2^S483A^ and Flag-RIOK2^S483D^ were performed with different sets of Flag-RIOK2^WT^ as control. A representative WB experiment for Flag-RIOK2^WT^ is shown. *(B)* Quantification of released RIOK2 to bound RIOK2 at t0 WB signal ratios from *(A)* using ImageLab software, n=3. *(C)* RIOK2^WT^ and RIOK2^S483A^ eHAP1 cells were treated with Leptomycin B (LMB, 20 nM) for the indicated times. Subcellular localization of RIOK2 and NOB1 was monitored by immunofluorescence microscopy using specific antibodies. Nuclei were visualized by DAPI staining. *(D), (E)*, Quantification of nuclear to cytoplasmic fluorescence ratios at the indicated time points obtained in *(C)* for RIOK2 *(D)* and NOB1 *(E)* using ImageJ software (n=100 cells from different fields). Statistically significant differences are indicated by asterisks (***: P≤0.001, *: P≤0.05, 2way ANOVA tests, Bonferroni posttests).

We anticipated that defects in RIOK2 dissociation from cytoplasmic pre-40S particles would slow down its recycling to the nucleus. To investigate the dynamics of RIOK2 shuttling, we compared the kinetics of RIOK2^WT^ or RIOK2^S483A^ nuclear import upon inhibition of pre-40S particle export using leptomycin B (LMB), an inhibitor of the CRM1 exportin. RIOK2^WT^ or RIOK2^S483A^ eHAP1 cell lines were treated with LMB and the nucleo-cytoplasmic distribution of RIOK2 was analyzed by IF during a time-course of LMB treatment (Fig. 6C), and quantified (Fig. 6D and 6E). In cells expressing RIOK2^WT^, the vast majority of cytoplasmic RIOK2 was imported back to the nucleus by 60 min of LMB treatment and the cytoplasmic signal became barely detectable by 90 min (Fig. 6C). In sharp contrast, RIOK2^S483A^ remained mostly cytoplasmic after 60 minutes of LMB treatment and a stronger signal remained in the cytoplasm after 90 min compared to RIOK2^WT^. Quantification of the nucleo-cytoplasmic ratios of the IF signals clearly confirmed the slower nuclear import rate of RIOK2^S483A^ compared to RIOK2^WT^ (Fig. 6D). We obtained similar results for NOB1, whose nuclear import following LMB treatment occurred more slowly in cells expressing RIOK2^S483A^ compared to control cells (Fig. 6C, lower panels and Fig. 6E). We concluded that RIOK2 phosphorylation at Ser483 facilitates its release and that of other AMFs from cytoplasmic pre-40S and allows their recycling into the nucleus.

## Discussion

The MAPK signaling pathway ensures coordinated expression of ribosome components and of the machinery involved in pre-ribosome assembly and maturation (Kusnadi et al. 2015). Upon activation, the MAPK pathway promotes both rDNA transcription by Pol I and Pol III in the nucleus, and translation of mRNAs encoding RPs and AMFs in the cytoplasm. Our study provides solid evidence that MAPK signaling applies another level of coordination during ribosome biogenesis, by directly regulating pre-ribosome assembly and maturation. RSK inhibition induces processing defects at different stages of the maturation process, suggesting that RSK regulates multiple steps of pre-rRNA processing. In agreement with this, our *in silico* search for canonical RXRXXS/T phosphorylation motifs identified potential phosphorylation sites in several other AMFs besides RIOK2.

We report direct evidence showing that RSK stimulates the maturation of pre-40S particles by facilitating the release of RIOK2 and other AMFs, thereby allowing the subsequent maturation steps to proceed towards completion of small subunit biogenesis (Fig. 7). We identified RIOK2 as a direct phosphorylation target of RSK and used a combination of approaches to gain in-depth functional understanding of how phosphorylation by RSK influences RIOK2 function.

**Figure 7.**
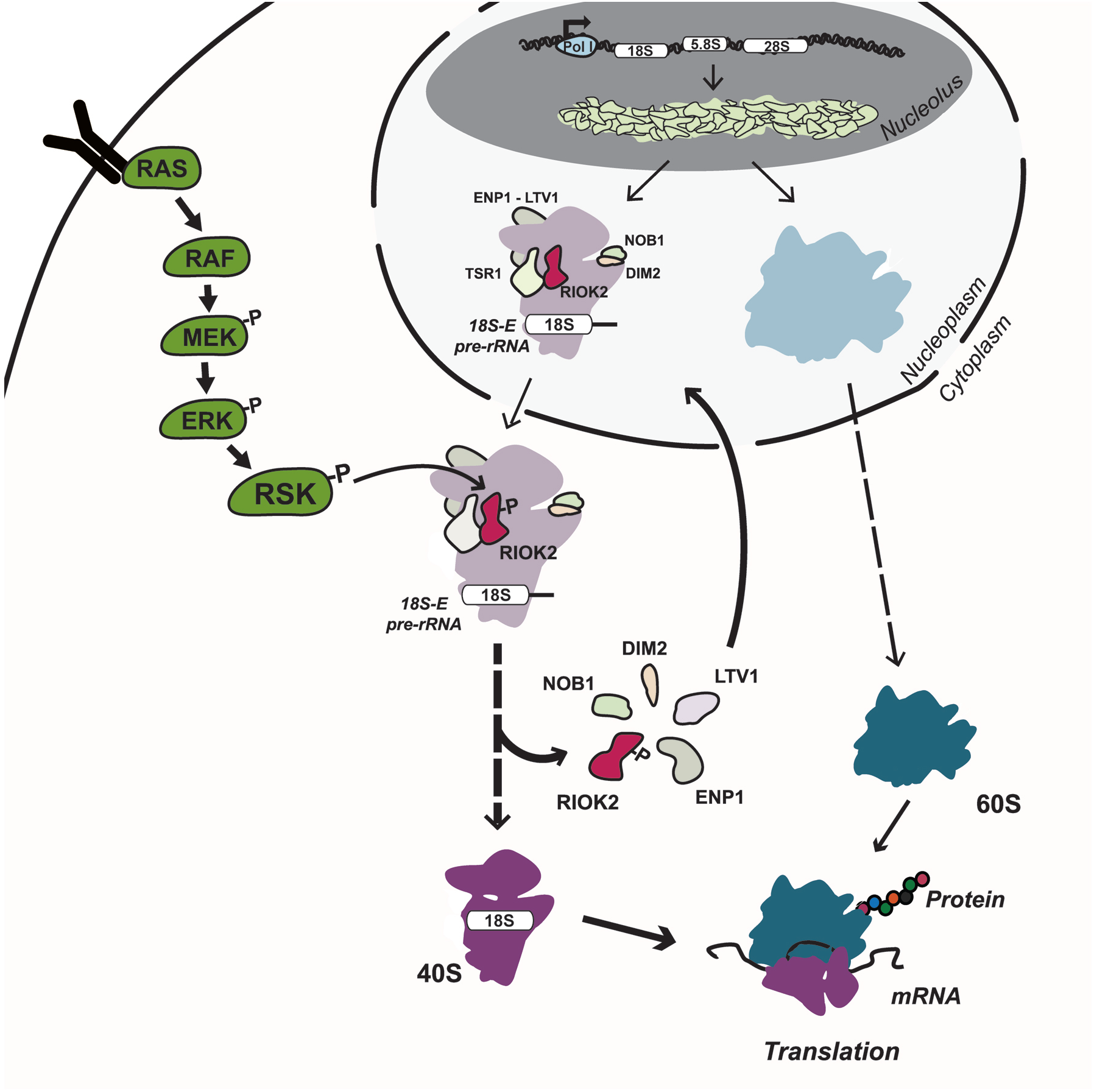
RIOK2 phosphorylation at Ser483 by RSK facilitates late stages of pre-40S particle maturation. RIOK2 is incorporated into pre-40S particles in the nucleus and participates to their export to the cytoplasm. Phosphorylation of RIOK2 by RSK at Ser483 facilitates its dissociation from pre-40S particles, which allows simultaneous or subsequent dissociation of other factors (ENP1, LTV1, DIM2, NOB1) to promote efficient maturation of the 18S-E pre-rRNA. This phosphorylation event is required for optimal protein synthesis and cell proliferation.

We demonstrated that RIOK2 phosphorylation at Ser483 stimulates the maturation of the 18S-E pre-rRNA by facilitating its release from pre-40S particles and promoting its recycling to the nucleus. It remains unclear whether RSK phosphorylates RIOK2 within pre-40S particles or before its association. A recent cryo-electron microscopy (cryo-EM) structure of a late human pre-40S particle containing RIOK2 has been reported (Ameismeier et al. 2018). The domain of RIOK2 containing Ser483 is not resolved in the structure (from V300 to C493), suggesting that this domain is highly flexible. Given that the flanking residues (E299 and S494) are located on the external surface of the pre-40S particle, it seems that this domain is protruding outside of the protein, and would therefore be accessible to RSK kinase for phosphorylation (Supplemental Fig. S5). These observations suggest that RSK could phosphorylate RIOK2 once incorporated into pre-40S particles to stimulate its dissociation. RSK has not been detected in pre-40S particles purified using different baits (Wyler et al. 2011), suggesting that its interaction with pre-40S particles may be very transient or labile. Consistent with this, sucrose gradient experiments show that RSK can be detected to low levels in the fractions containing pre-40S particles (Supplemental Fig. S6).

The molecular mechanisms underlying yeast Rio2 release from pre-40S particles have been recently investigated (Knüppel et al. 2018). Rio2 binds to pre-40S particles in a catalytically inactive conformation with its catalytic P-loop lysine (Lys106) bound to the pre-rRNA (Ferreira-Cerca et al. 2012). Following unclear conformational rearrangements within the pre-40S particles, release of the P-loop Lys106 allows Rio2 activation and dissociation from the particles. In the human pre-40S particle, RIOK2 seems to be positioned in a similar way to the yeast particle, with its P-loop Lys105 in close contact to helix 30 of the 18S rRNA (Supplemental Fig. S5). However, as the C-terminal extension of RIOK2 containing Ser483 is not present in yeast Rio2, some aspects of the function or regulation of the protein are likely different in yeast and human cells. We propose that phosphorylation of RIOK2 at Ser483 by RSK could participate in conformational rearrangements that trigger RIOK2 catalytic activity by releasing its P-loop lysine and/or weakening its association with pre-40S particles, both favoring RIOK2 dissociation from pre-40S particles.

Part of the data supporting our conclusion that RIOK2 phosphorylation at Ser483 is required for optimal maturation of pre-40S particles stems from our *in vivo* experiments where RIOK2 Ser483 was substituted to an alanine using the CRISPR/Cas9 approach. We cannot exclude that what is important for efficient 18S-E pre-rRNA processing is Ser483 *per se*, and not its phosphorylation by RSK. However, we consider this hypothesis unlikely for several reasons. Substitution of Ser483 to an aspartic acid instead of an alanine (RIOK2^S483D^ mutant) does not induce an accumulation of 18S-E pre-rRNA (Fig. 4C and 4D), indicating that the serine residue can be substituted without significant consequences by a residue mimicking the phosphorylated state. In our i*n vitro* dissociation assay, RIOK2^S483D^ behaves like the wild-type protein, indicating again that a negative charge at position 483 is sufficient to confer close to wild-type physical properties to RIOK2. Finally, release of assembly factors by phosphorylation events is not unprecedented. In yeast, Hrr25 kinase phosphorylates Rps3 and Enp1 within nuclear pre-40S particles and thereby weakens their association with the particle to promote conformational rearrangements necessary for formation of the beak structure (Schäfer et al. 2006). Hrr25/CK1 also promotes the dissociation of Ltv1 from pre-40S particles both in human and yeast (Ghalei et al. 2015; Zemp et al. 2014). These data combined with our study suggest that phosphorylation events are a common theme in the regulation of AMF dissociation from pre-ribosomal particles.

Inhibition of RSK-dependent RIOK2 phosphorylation, although it delays 18S-E pre-rRNA processing, does not affect the steady-state amount of ribosomes (18S rRNA levels). We do observe, however, a global defect in translation correlated with a decrease in cell proliferation. Several hypotheses can be proposed to account for this apparent paradox. For technical reasons, we may have been unable to detect minor reductions in mature 18S rRNA levels. Another hypothesis could be that cells may extend the half-life of the ribosomes produced in these conditions to counterbalance a production defect. These ribosomes may become partially dysfunctional due to increased exposure to reactive oxygen species, leading to stalled ribosomes or decreased translation efficiency (Mills and Green 2017).

RIOK2 is the first pre-ribosome AMF shown to be regulated by the MAPK pathway. Since this process involves hundreds of AMFs, potentially many other MAPK-driven regulatory mechanisms operate. Like RIOK2, a significant proportion of AMFs include energy-consuming enzymes such as kinases, ATPases, GTPases or RNA helicases (Henras et al. 2015; Aubert et al. 2018; Cerezo et al. 2019). These enzymes have been suggested to provide directionality, accuracy and quality control to the process. They are believed to provide energy to overcome thermodynamically unfavorable steps of the process such as disruption of stable RNA helices, protein/RNA or protein/protein interactions (Kressler et al. 2010; Strunk and Karbstein 2009). Our *in silico* screen identified several other potential substrates of RSK and among these, energy-consuming enzymes are particularly represented. We therefore propose that MAPK signaling may participate in the coordination of the series of events occurring during pre-ribosome assembly and maturation by stimulating the activity of selected energy-consuming enzymes, thereby allowing to overcome rate-limiting steps. Interestingly, mammalian RIOK1 and RIOK3, also involved in pre-40S particle maturation (Baumas et al. 2012; Widmann et al. 2012) are not phosphorylated within a RSK consensus motif (Baumas et al. 2012; Widmann et al. 2012) suggesting a specific regulation of RIOK2 by RSK.

Our study paves the way for future exploration of the regulation of ribosome biogenesis at the post-transcriptional level by the MAPK pathway. Identification of key limiting steps, in particular those catalyzed by discrete catalytic activities, would help in designing innovative molecules aimed at counteracting MAPK-driven deregulated ribosome production in pathologies, such as cancers or RASopathies.

## Materials and Methods

### Cloning

Plasmids and sequences of oligonucleotides used in this study are listed in Supplemental Table 2. All clonings have been performed using In-Fusion HD Cloning Plus (Takara, Cat#638911) according to manufacturer’s recommendations. Transformations have been performed using Stellar(tm) Competent Cells (Takara, Cat#636763).

### Human cell lines, transfections and chemicals

All human cell lines used in this study are listed in Reporting Summary. Human cells were maintained in 5% CO_2_ at 37°C. HEK293 and HeLa cells were cultured in Dulbecco’s modified Eagle’s medium (DMEM) and eHAP1 cells in Iscove Modified Dulbecco Media (IMDM). Both media were supplemented with 10% Fetal Bovine Serum, 1% Penicillin-Streptomycin, 1% Pyruvate. When indicated, cells were treated with Phorbol 12-Myristate 13-Acétate (Fisher Scientific, Cat#10061403), Human EGF (Euromedex, Cat# HC88823), LJH685 (Selleck Chemicals, S7870), BI-D1870 (Selleck Chemicals, S2843) and/or PD184352 (Selleck Chemicals, S1020). For transient plasmid expression, cells were transfected using either Jet Prime reagent or calcium phosphate precipitation. For shRNA-mediated RSK1/2 knockdown, cells were infected by lentiviruses produced with vectors from the Mission TRC shRNA library (RSK1, TRCN470; RSK2, TRCN537) in the presence of 4 mg/ml polybrene and selected 48 h after infection with 2 µg/mL puromycin.

### CRISPR/Cas9 genome editing

RIOK2^S483A^ and RIOK2^S483D^ eHAP1 mutant cell lines were generated using CRISPR/Cas9-mediated double strand break and homologous recombination, using ouabain co-selection as described (Agudelo et al. 2017). Oligos designed to encode the Cas9 guide RNA (http://crispor.tefor.net/) were annealed and ligated into BbsI-digested Addgene 86613 plasmid, resulting in plasmid 86613-RIOK2-gRNA. Single-stranded donor templates were designed to introduce the RIOK2^S483A^ or RIOK2^S483D^ point mutations along with silent mutations introducing an MscI or EcoRV restriction site, respectively. Plasmid 86613-RIOK2-gRNA and single-stranded donor templates for introduction of ouabain resistance and either RIOK2^S483A^ or RIOK2^S483D^ point mutations were electroporated into eHAP1 cells at 300 V with a Gene Pulser System (Bio-Rad Laboratories) in cuvettes with a 4-mm inter-electrode distance (Eurogentec). Transfected cells were grown for 48 h in the presence of 7 µM ouabain (Sigma, O3125; CAS:11018-89-6), then diluted into 14 cm-dishes and grown for 2-3 weeks in the presence of ouabain to obtain isolated clones. Clonal populations were isolated into 12 well-plates. To identify mutant cells, genomic DNAs were isolated from clonal populations using GenElute(tm) Mammalian Genomic DNA Miniprep Kit Protocol (Sigma-Aldrich). A genomic region of RIOK2 gene encompassing Ser483 was PCR-amplified from the genomic DNAs, and the presence of point mutations was revealed by digestion of PCR products with MscI (for RIOK2^S483A^) or EcoRV (for RIOK2^S483D^). Homozygous knock in clones were then confirmed by sequencing (Eurofins Genomics). The wild-type controls used in the study are randomly chosen eHAP1 cell lines electroporated with 86613-RIOK2-gRNA plasmid and donor templates but in which RIOK2 locus had not been edited.

### Proliferation and cell death analyses

Cell proliferation was assessed using either MTS assay (CellTiter 96® AQueous One Solution Cell Proliferation Assay, Promega, data are expressed as means of three repeated measures using at least 5 distinct samples for each condition) or cell counting (data are expressed as means of three repeated measures using 3 distinct samples for each condition). Cell death was monitored using FITC Annexin V Apoptosis Detection Kit (BioLegend). Populations of cells in apoptosis, necrosis or post-apoptosis were discriminated using a FACS Verse analyzer (BD Biosciences, data are expressed as means of 3 distinct samples for each condition).

### Protein Synthesis assay

Global protein synthesis was determined using Sunset method (Schmidt et al. 2009). Puromycin (InVivoGen, Cat#ant-pr; CAS: 58-58-2) was added in the culture medium (1µM) and cells were incubated for 20, 30 or 40 min at 37°C. After cell lysis, normalized amounts of total proteins were analyzed by Western Blot using anti-puromycin antibodies.

### RNA analyses

Extractions of total RNAs were performed using TRI REAGENT (MRC). After addition of 0.3 mL of cholorophorm per mL of TRI REAGENT, the mixtures were shaken vigorously and centrifuged at 12,000 g for 15 minutes at 4°C. Aqueous phases were submitted to a second round of extraction using 0.5 volume or water-saturated phenol and 0.5 volume of chloroform. For precipitation, RNAs were mixed with one volume of 2-propanol and incubated 10 min at room temperature. RNAs were pelleted by centrifugation at 12,000 × g for 10 minutes at 2–8 °C. RNA pellets were washed with 1 mL of 75% ethanol and centrifugation at 12,000 × g for 10 minutes at 2–8 °C. Air-dried pellets were resuspended with ultrapure MilliQ H_2_O and RNAs were quantified using a NanoDrop Spectrophotometer (Thermo Fisher Scientific). Northern blot experiments were performed as described in “Molecular Cloning”, Sambrook and Russell, CSHL Press (“Separation of RNA According to Size: Electrophoresis of Glyoxylated RNA through Agarose Gels”). Briefly, equal amounts of total RNAs (usually 4 µg for analysis of rRNA precursors or 1 µg for mature rRNAs) were mixed with five volumes of Glyoxal loading buffer [prepared by mixing the following: 6 ml DMSO, 2 ml deionized glyoxal, 1.2 ml 10X BPTE (see below), 600 µl 80% glycerol, 40 µl 10 mg/ml Ethidium Bromide]. The samples were heated 1 hour at 55°C and RNAs were separated by electrophoresis on 1.2 % agarose gels in 1X BPTE running buffer [100 mM PIPES, 300 mM BIS-TRIS, 10 mM EDTA]. After electrophoresis, the gels were (i) rinsed 2 times 5 min with ultrapure MilliQ H_2_O, (ii) soaked 20 min at room temperature (RT) in 75 mM NaOH with gentle shaking to partially hydrolyze RNAs, (iii) rinsed 2 times 5 min with ultrapure MilliQ H_2_O, (iv) soaked 2 times 15 min at RT in [0.5 M Tris-HCl pH 7.4, 1.5 M NaCl] with gentle shaking to neutralize the pH, (v) Soaked 2 times 10 min at RT in 10X SSC with gentle shaking. RNAs were then transferred over night to Amersham Hybond N^+^ membranes (GE Healthcare) by capillarity with 10X SSC transfer buffer. Membranes were then exposed to 0.125 joules of 365 nm UV rays to crosslink RNAs on the membranes. Membranes were then hybridized with ^32^P-labeled oligonucleotide probes using the Rapid-hyb buffer (GE Healthcare). Radioactive membranes were exposed to PhosphorImager screens and signals were revealed using Typhoon imager (GE Healthcare). The sequences of the probes used to detect (pre-)rRNAs are described in Supplemental Table 2. Quantification of RNA levels was obtained using MultiGauge software (Fujifilm).

### Protein analyses

Protein extracts were prepared as follow: cells were washed with ice-cold PBS, and lysed with Buffer A [10 mM K_3_PO_4_, 1 mM EDTA, 5 mM EGTA, 10 mM MgCl_2_, 50 mM β-glycerophosphate, 0.5 % Nonidet P-40, 0.1 % Brij 35, cOmplete protease inhibitor cocktail (Roche), Phosphatase Inhibitor Cocktail 2 and 3 (Sigma-Aldrich)]. Protein concentrations were measured using Bio-Rad Protein Assay and normalized protein concentrations were resuspended in Laemmli Buffer [40 mM Trizma base, 2 % SDS, 5 % Glycerol, 0.08 % Bromophenol blue, 25 mM DTT]. Western blot experiments were performed as follow: protein samples were heated 5 min at 95°C, loaded on SDS-polyacrylamide gels (8 to 10 %) and transferred to nitrocellulose membranes using Trans-blot turbo transfer system (Bio-Rad). Membranes were saturated for 1 h with TBST buffer (150 mM NaCl, 20 mM Tris pH 8.0, 0.001% Tween-20) containing 5% powder milk, and incubated over night with the same buffer containing primary antibodies. After 3 washes with TBST buffer, membranes were incubated for 1 h with the secondary antibodies diluted in TBST containing 5% powder milk, and washed three times with TBST buffer before ECL detection. ImageLab software (Biorad) was used to quantify protein signals from Western Blot.

### In-gel tryptic digestion and nanoLC-MS/MS analysis

For mass spectrometry analysis, HA-RIOK2 immunoprecipitated samples, prepared in triple biological replicates for each condition, were reduced for 30 min at 55°C in Laemmli buffer containing 25 mM DTT and alkylated in 90 mM iodoacetamide for 30 min in the dark at room temperature. Equal volumes of samples were separated by SDS–PAGE on 10% polyacrylamide gels, followed by gel staining with InstantBlue™ (Expedeon Protein Solutions) according to the manufacturer’s instructions. Bands at the molecular weight of HA-RIOK2 were excised and subjected to in-gel tryptic digestion using modified porcine trypsin (Promega) at 20 ng/μl as previously described (Shevchenko et al., 1996). The dried peptide extracts obtained were resuspended in 21 µl of 0.05% trifluoroacetic acid in 2% acetonitrile spiked-in with 0.1X final concentration of iRT standard peptides (Biognosis) and analyzed by online nanoLC using UltiMate 3000 RSLCnano LC system (ThermoScientific, Dionex) coupled to an Orbitrap Fusion Tribrid mass spectrometer (Thermo Scientific, Bremen, Germany). 5µl of each peptide extracts were loaded onto 300 µm ID x 5 mm PepMap C18 precolumn (ThermoFisher, Dionex) at 20 µl/min in 2% acetonitrile, 0.05% trifluoroacetic acid. After 5 min of desalting, peptides were online separated on a 75 µm ID x 50 cm C18 column (in-house packed with Reprosil C18-AQ Pur 3 μm resin, Dr. Maisch ; Proxeon Biosystems, Odense, Denmark), equilibrated in 95% of buffer A (0.2% formic acid), with a gradient of 5 to 25% of buffer B (80% acetonitrile, 0.2% formic acid) for 80 min then 25% to 50% for 30 min at a flow rate of 300 nl/min. The instrument was operated in the data-dependent acquisition (DDA) mode using a top-speed approach (cycle time of 3 s). The survey scans MS were performed in the Orbitrap over m/z 350–1550 with a resolution of 120,000 (at 200 m/z), an automatic gain control (AGC) target value of 4e5, and a maximum injection time of 50 ms. Most intense ions per survey scan were selected at 1.6 m/z with the quadrupole and fragmented by Higher Energy Collisional Dissociation (HCD). The monoisotopic precursor selection was turned on, the intensity threshold for fragmentation was set to 50,000 and the normalized collision energy was set to 35%. The resulting fragments were analyzed in the Orbitrap with a resolution of 30,000 (at 200 m/z), an automatic gain control (AGC) target value of 5e4, and a maximum injection time of 60 ms. The dynamic exclusion duration was set to 30 s with a 10 ppm tolerance around the selected precursor and its isotopes. For internal calibration the 445.120025 ion was used as lock mass. Triplicate technical LC-MS measurements were performed for each sample.

### Database search and label-free quantitative analysis of RIOK2 phosphorylation

All raw MS files were processed with MaxQuant (v 1.5.2.8) for database search with the Andromeda search engine and quantitative analysis. Data were searched against the UniProtKB/Swiss-Prot protein database released 2015_07 with *Homo sapiens* taxonomy (11953 sequences) supplemented with the human 3HA-RIOK2 sequence, the Biognosys iRT peptide sequences and a list of frequently observed contaminant sequences provided in MaxQuant 1.5.2.8. Carbamidomethylation of cysteines was set as a fixed modification, whereas oxidation of methionine, protein N-terminal acetylation, and phosphorylation of serine, threonine, and tyrosine were set as variable modifications. Enzyme specificity was set to trypsin/P, and a maximum of three missed cleavages was allowed. The precursor mass tolerance was set to 20 ppm for the first search and 10 ppm for the main Andromeda database search, and the mass tolerance in MS/MS mode was set to 0.025 Da. The required minimum peptide length was seven amino acids, and the minimum number of unique peptides was set to one. Andromeda results were validated by the target-decoy approach using a reverse database and the false discovery rates at the peptide-spectrum matches (PSM), protein and site levels were set to 1%. Phosphosite localization was evaluated on the basis of the Phosphosite Localization Scoring and Localization Probability algorithm of the Andromeda search engine. For label-free relative quantification of the samples, the match between runs option of MaxQuant was enabled with a time window of 2 min, to allow cross-assignment of MS features detected in the different runs. Relative quantification of RIOK2 phosphorylation sites was performed by retrieving the intensity values of the phosphorylated peptide ions from the MaxQuant evidence.txt output that contains quantitative data for all peptide ions. Intensity values were first normalized for instrument variation using the MS intensities of the iRT spiked-in standards. The variability that may occur during the immunopurification was then corrected in each sample by normalizing the iRT-normalized intensity values to that of the HA-RIOK2 bait. This second normalization was performed based on the sum of the intensity values of HA-RIOK2 tryptic peptides and, to exclude variations resulting from RIOK2 phosphorylation, all of the HA-RIOK2 peptides containing a residue susceptible to phosphorylation were eliminated from the calculation. Mean intensity values were then calculated from technical LC-MS replicates. To evaluate the relative abundance of phosphorylation at a given site, total areas of tryptic peptides encompassing the site were calculated for the phosphorylated forms by aggregating data corresponding to peptide ions charge states (2+ and 3+), modification other than phosphorylation (oxidized methionine), and tryptic miscleavages (overlapping sequences).

### Immunoprecipitations

HEK293 cells (one 10 cm-dish at 90% confluence per condition) were lysed in buffer A supplemented with 0.1 M KCl. Cell lysates were incubated with the indicated antibodies for 1 h 45 min, then with protein A-Sepharose CL-4B beads (GE Healthcare) for another 45 min. Immunoprecipitates were washed 3 times with lysis buffer and eluted from the beads upon addition of Laemmli buffer and incubation 5 min at 95°C.

### In vitro dissociation assay

Pre-40S particles were immunoprecipitated as described above from HEK293 cells expressing Flag-RIOK2 and HA-NOB1 (one 10 cm-dish at 90% confluence per condition). After the third wash with buffer A supplemented with 0.1M KCl, beads were washed once with “RIOK2 dissociation buffer” (200 mM NaCl, 25 mM Tris-HCl pH 7,4, 10 mM MgCl_2_, 5 mM β-glycerophosphate). HA-NOB1-associated pre-40S particles were then incubated for indicated times in “RIOK2 dissociation buffer” supplemented with 1mM ATP at 16°C. Beads and supernatants were then separated and mixed with Laemmli buffer.

### Sucrose gradient fractionation

Three days after seeding, culture medium of two 15 cm-dishes of HEK293 cells at 90% confluence was removed and fresh 37°C-prewarmed medium was added to the cells. After an incubation of ∼90 min at 37°C, 10 µg/ml cycloheximide was added directly to the culture medium and incubation was prolonged for 10 min. Cells were harvested with trypsin and washed 2 times with ice-cold PBS supplemented with 10 µg/ml cycloheximide. The cell pellet was then washed with buffer B (10 mM HEPES–KOH pH 7.9, 10 mM KCl, 1.5 mM MgCl2, 100 µg/ml cycloheximide) and incubated 20 min on ice in buffer B supplemented with 0.5 mM dithiothreitol, 1 × cOmplete EDTA-free protease inhibitor cocktail (Roche) and 0.5 U/µl RNasin (Promega). After incubation cells were disrupted using a Dounce homogenizer with a tight pestle and centrifuged at 1000 x g for 10 min at 4°C, in order to pellet nuclei. The top soluble phase, containing the cytoplasmic fraction, was clarified through one centrifugation at 10 000 x g for 15 min at 4°C and quantified by measuring absorbance at 260 nm. Normalized amounts of extracts were loaded on a 10–50% sucrose gradient in buffer B. Gradients were centrifuged at 39 000 rpm for 2.5 h at 4°C in an Optima L-100XP ultracentrifuge (Beckman– Coulter) using the SW41Ti rotor with brake. Following centrifugation, the fractions were collected using a Foxy Jr fraction collector (Teledyne ISCO) and the absorbance at 254 nm was measured with a UA-6 device (Teledyne ISCO). For protein analyses, fractions were precipitated with TCA and protein pellets were resuspended in Laemmli buffer.

### Microscopy

For all microscopy experiments, cells were seeded on microscope cover glasses in 6-well plates and grown for 48 to 72 h. Immunofluorescence (IF) microscopy experiments were performed as described previously (Zemp et al. 2009). Briefly, after fixation in 4% paraformaldehyde (PFA), cells were permeabilized with [0.1% Triton X-100 and 0.02% SDS in PBS] for 5 min. Fixed cells were incubated in blocking solution [2% BSA (Sigma A8022) in PBS] for 30 min and then incubated for 1h with the same solution containing primary antibodies diluted to 1:2000. Cells were washed 3 times for 5 min with [2% BSA in PBS], and subsequently incubated for 30 min with secondary antibodies (Alexa Fluor 488-conjugated goat anti-rabbit antibodies) diluted in blocking solution. After 3 washes, cells were incubated briefly in [0.1% Triton X-100, 0.02% SDS in PBS], and post-fixed with 4% PFA. After a wash with PBS, coverslip were mounted in VectaShield (Vector Laboratories). Fluorescent *In Situ* Hybridization (FISH) experiments were done as described previously (O’Donohue et al. 2010). Cells were fixed in 4% PFA, and after 2 washes with PBS, cells were permeabilized at 4°C for 18 h in 70% ethanol. Permeabilized cells were washed twice in (2X SSC, 10% formamide) and hybridized at 37°C in the dark for ≥ 5 h in hybridization buffer (10% formamide, 0.1X SSC, 0.5 mg/ml *E. coli* tRNAs, 10% dextran sulfate, 250 μg/ml BSA, 10 mM ribonucleoside vanadyl complexes, 0.5 ng/μL of Cy3-conjugated 5’ITS1 probe). After 2 washes in (2X SSC, 10% formamide), cells were rinsed with PBS, and coverslip were mounted in VectaShield. Images were captured using an inverted Olympus IX81 epifluorescence microscope equipped with a X100 objective lens (UPlan SApo 1.4 oil), a SpectraX illumination system (Lumencore©) and a CMOS camera (Hamamatsu© ORCA-Flash 4.0), driven by MetaMorph (Molecular Devices). Fluorescent signals were captured after different exposure times (between 500 and 2000 ms) depending on signal intensities. Image analyses were performed using ImageJ software.

### In vitro kinase assay

For RSK kinase assays, recombinant-activated RSK1 purchased from SignalChem was used with bacterially purified recombinant GST-RIOK2 (aa 443–552) as substrate (WT and S483A), under linear assay conditions. Assays were performed for 10 min at 30°C in kinase buffer [25 mmol/L Tris-HCl (pH 7.4), 10 mmol/L MgCl_2_, and 5 mmol/L β-glycerophosphate] supplemented with 5 μCi of [γ-^32^P]ATP. All samples were subjected to SDS-PAGE followed by immunoblotting, and incorporation of radioactive ^32^P label was determined by autoradiography using a Fuji PhosphorImager with ImageQuant software. The data presented are representative of at least three independent experiments.

### Statistical Analyses

Data are expressed as means ± SEM. All statistical data (n≥3) were calculated using GraphPad Prism 5.01. Statistical details and significance reports can be found in the corresponding figure legends.

## Acknowledgments

We are grateful to Christian Montelese, Ivo Zemp and Ulrike Kutay (ETH Zurich) for sharing antibodies, technical advice and helpful scientific discussions. Our work as benefited from fruitful technical and scientific contacts with the neighboring groups at LBME/CBI of P.E. Gleizes (MF. O’Donohue, C. Plisson, M. Aubert), J. Cavaillé (J. Hebras), K. Bystricky (L. Recoules) and T. Kiss (S. Egloff, P. Vitali). Mass spectrometry analyses were performed at the IPBS proteomic facility led by Odile Burlet-Schiltz in collaboration with Carine Froment. Fluorescence microscopy was performed at the LITC (Light Imaging Toulouse CBI) facility with the expertize of Sylvain Cantaloube. FACS analyses were performed at the Cytology and Cell Sorting facility at the I2MC, Toulouse with the help of Elodie Riant. We thank Dominique Helmlinger (CRBM, Montpellier) and Hervé Prats (CRCT, Toulouse) for their advice and scientific input to the study. We thank Deborah Carper for advice with statistics. We thank all members of the Henry/Henras team for helpful discussions. The Henry/Henras group is supported by grants from ANR (to A.H.) and Ligue Contre le Cancer (to Y.R.). The work was also supported in part by the Région Midi-Pyrénées, European funds (Fonds Européens de Développement Régional, FEDER), Toulouse Métropole, and by the French Ministry of Research with the Investissement d’Avenir Infrastructures Nationales en Biologie et Santé program (ProFI, Proteomics French Infrastructure project, ANR-10-INBS-08). This work was also supported by grants from the Canadian Institutes for Health Research (MOP-142374 and PJT-152995 to P.P.R.), and the Natural Sciences and Engineering Research Council of Canada (P.P.R.). P.P.R. is a scholar of the Fonds de la recherche du Québec - Santé (FRQS). E.C. is supported by a fellowship from the French Ministry of Higher Education and Research.

## Author contributions

E.L.C., A.K.H. and Y.R. designed all experiments, with help from Y.H. and P.P.R. All experiments were performed by E.L.C., T.H., G.L., C.F., A.K.H. and Y.R., with help from S.C., O.L., M.-K.S., C.A. and M.H. All authors interpreted the data, and E.L.C., Y.R. and A.K.H. wrote the manuscript with contributions from Y.H. and P.P.R.

## References

1. Henras, A. K., Plisson-Chastang, C., O’Donohue, M.-F., Chakraborty, A. & Gleizes, P.-E. An overview of pre-ribosomal RNA processing in eukaryotes. Wiley Interdiscip Rev RNA 6, 225–242 (2015).

2. Kressler, D., Hurt, E. & Baßler, J. A puzzle of life: crafting ribosomal subunits. Trends Biochem. Sci. 42, 640–654 (2017).

3. Aubert, M., O’Donohue, M.-F., Lebaron, S. & Gleizes, P.-E. Pre-Ribosomal RNA Processing in Human Cells: From Mechanisms to Congenital Diseases. Biomolecules 8, (2018).

4. Cerezo, E. et al. Maturation of pre-40S particles in yeast and humans. Wiley Interdiscip Rev RNA 10, e1516 (2019).

5. Woolford, J. L. & Baserga, S. J. Ribosome biogenesis in the yeast Saccharomyces cerevisiae. Genetics 195, 643–681 (2013).

6. Wild, T. et al. A protein inventory of human ribosome biogenesis reveals an essential function of exportin 5 in 60S subunit export. PLoS Biol. 8, e1000522 (2010).

7. Tafforeau, L. et al. The complexity of human ribosome biogenesis revealed by systematic nucleolar screening of Pre-rRNA processing factors. Mol. Cell 51, 539–551 (2013).

8. Badertscher, L. et al. Genome-wide RNAi Screening Identifies Protein Modules Required for 40S Subunit Synthesis in Human Cells. Cell Rep. 13, 2879–2891 (2015).

9. Farley-Barnes, K. I. et al. Diverse regulators of human ribosome biogenesis discovered by changes in nucleolar number. Cell Rep. 22, 1923–1934 (2018).

10. Bohnsack, K. E. & Bohnsack, M. T. Uncovering the assembly pathway of human ribosomes and its emerging links to disease. EMBO J. 38, e100278 (2019).

11. Goodfellow, S. J. & White, R. J. Regulation of RNA polymerase III transcription during mammalian cell growth. Cell Cycle 6, 2323–2326 (2007).

12. Grummt, I. Life on a planet of its own: regulation of RNA polymerase I transcription in the nucleolus. Genes Dev. 17, 1691–1702 (2003).

13. Mahoney, S. J., Dempsey, J. M. & Blenis, J. in Translational Control in Health and Disease 90, 53–107 (Elsevier, 2009).

14. Piazzi, M., Bavelloni, A., Gallo, A., Faenza, I. & Blalock, W. L. Signal transduction in ribosome biogenesis: A recipe to avoid disaster. Int. J. Mol. Sci. 20, (2019).

15. Cargnello, M. & Roux, P. P. Activation and function of the MAPKs and their substrates, the MAPK-activated protein kinases. Microbiol. Mol. Biol. Rev. 75, 50–83 (2011).

16. Houles, T. & Roux, P. P. Defining the role of the RSK isoforms in cancer. Semin. Cancer Biol. 48, 53–61 (2018).

17. Romeo, Y., Zhang, X. & Roux, P. P. Regulation and function of the RSK family of protein kinases. Biochem. J. 441, 553–569 (2012).

18. Wullschleger, S., Loewith, R. & Hall, M. N. TOR signaling in growth and metabolism. Cell 124, 471–484 (2006).

19. Roux, P. P. & Topisirovic, I. Signaling Pathways Involved in the Regulation of mRNA Translation. Mol. Cell. Biol. 38, (2018).

20. Kusnadi, E. P. et al. Regulation of rDNA transcription in response to growth factors, nutrients and energy. Gene 556, 27–34 (2015).

21. Zhao, J., Yuan, X., Frödin, M. & Grummt, I. ERK-dependent phosphorylation of the transcription initiation factor TIF-IA is required for RNA polymerase I transcription and cell growth. Mol. Cell 11, 405–413 (2003).

22. Felton-Edkins, Z. A. et al. The mitogen-activated protein (MAP) kinase ERK induces tRNA synthesis by phosphorylating TFIIIB. EMBO J. 22, 2422–2432 (2003).

23. Sriskanthadevan-Pirahas, S., Deshpande, R., Lee, B. & Grewal, S. S. Ras/ERK-signalling promotes tRNA synthesis and growth via the RNA polymerase III repressor Maf1 in Drosophila. PLoS Genet. 14, e1007202 (2018).

24. Stefanovsky, V. Y. et al. An immediate response of ribosomal transcription to growth factor stimulation in mammals is mediated by ERK phosphorylation of UBF. Mol. Cell 8, 1063–1073 (2001).

25. Stefanovsky, V., Langlois, F., Gagnon-Kugler, T., Rothblum, L. I. & Moss, T. Growth factor signaling regulates elongation of RNA polymerase I transcription in mammals via UBF phosphorylation and r-chromatin remodeling. Mol. Cell 21, 629–639 (2006).

26. Stefanovsky, V. Y., Langlois, F., Bazett-Jones, D., Pelletier, G. & Moss, T. ERK modulates DNA bending and enhancesome structure by phosphorylating HMG1-boxes 1 and 2 of the RNA polymerase I transcription factor UBF. Biochemistry 45, 3626–3634 (2006).

27. Stefanovsky, V. Y. & Moss, T. The splice variants of UBF differentially regulate RNA polymerase I transcription elongation in response to ERK phosphorylation. Nucleic Acids Res. 36, 5093–5101 (2008).

28. Zhang, Z., Liu, R., Townsend, P. A. & Proud, C. G. p90(RSK)s mediate the activation of ribosomal RNA synthesis by the hypertrophic agonist phenylephrine in adult cardiomyocytes. J. Mol. Cell Cardiol. 59, 139–147 (2013).

29. Chauvin, C. et al. Ribosomal protein S6 kinase activity controls the ribosome biogenesis transcriptional program. Oncogene 33, 474–483 (2014).

30. Geyer, P. K., Meyuhas, O., Perry, R. P. & Johnson, L. F. Regulation of ribosomal protein mRNA content and translation in growth-stimulated mouse fibroblasts. Mol. Cell. Biol. 2, 685–693 (1982).

31. Asmal, M. et al. Production of ribosome components in effector CD4+ T cells is accelerated by TCR stimulation and coordinated by ERK-MAPK. Immunity 19, 535–548 (2003).

32. Romeo, Y. et al. RSK regulates activated BRAF signalling to mTORC1 and promotes melanoma growth. Oncogene 32, 2917–2926 (2013).

33. Iadevaia, V., Zhang, Z., Jan, E. & Proud, C. G. mTOR signaling regulates the processing of pre-rRNA in human cells. Nucleic Acids Res. 40, 2527–2539 (2012).

34. Isfort, R. J. & Cody, D. B. Serum and growth factors stimulate ribosomal RNA processing in Syrian hamster embryo cells: divergence of this signalling pathway from immediate-early gene expression. Cell Signal. 4, 665–674 (1992).

35. Vanrobays, E. et al. TOR regulates the subcellular distribution of DIM2, a KH domain protein required for cotranscriptional ribosome assembly and pre-40S ribosome export. RNA 14, 2061–2073 (2008).

36. Raman, N., Nayak, A. & Muller, S. mTOR signaling regulates nucleolar targeting of the SUMO-specific isopeptidase SENP3. Mol. Cell. Biol. 34, 4474–4484 (2014).

37. Talkish, J., Zhang, J., Jakovljevic, J., Horsey, E. W. & Woolford, J. L. Hierarchical recruitment into nascent ribosomes of assembly factors required for 27SB pre-rRNA processing in Saccharomyces cerevisiae. Nucleic Acids Res. 40, 8646–8661 (2012).

38. Honma, Y. et al. TOR regulates late steps of ribosome maturation in the nucleoplasm via Nog1 in response to nutrients. EMBO J. 25, 3832–3842 (2006).

39. LaRonde-LeBlanc, N. & Wlodawer, A. The RIO kinases: an atypical protein kinase family required for ribosome biogenesis and cell cycle progression. Biochim. Biophys. Acta 1754, 14–24 (2005).

40. Vanrobays, E., Gelugne, J.-P., Gleizes, P.-E. & Caizergues-Ferrer, M. Late cytoplasmic maturation of the small ribosomal subunit requires RIO proteins in Saccharomyces cerevisiae. Mol. Cell. Biol. 23, 2083–2095 (2003).

41. Asquith, C. R. M., East, M. P. & Zuercher, W. J. RIOK2: straddling the kinase/ATPase line. Nat. Rev. Drug Discov. 18, 574 (2019).

42. Read, R. D. et al. A kinome-wide RNAi screen in Drosophila Glia reveals that the RIO kinases mediate cell proliferation and survival through TORC2-Akt signaling in glioblastoma. PLoS Genet. 9, e1003253 (2013).

43. Song, Y. et al. RIOK2 is negatively regulated by miR-4744 and promotes glioma cell migration/invasion through epithelial-mesenchymal transition. J. Cell Mol. Med. (2020). doi:10.1111/jcmm.15107

44. Liu, T. et al. Phosphorylation of right open reading frame 2 (Rio2) protein kinase by polo-like kinase 1 regulates mitotic progression. J. Biol. Chem. 286, 36352–36360 (2011).

45. Zemp, I. et al. Distinct cytoplasmic maturation steps of 40S ribosomal subunit precursors require hRio2. J. Cell Biol. 185, 1167–1180 (2009).

46. Rouquette, J., Choesmel, V. & Gleizes, P.-E. Nuclear export and cytoplasmic processing of precursors to the 40S ribosomal subunits in mammalian cells. EMBO J. 24, 2862–2872 (2005).

47. Ferreira-Cerca, S. et al. ATPase-dependent role of the atypical kinase Rio2 on the evolving pre-40S ribosomal subunit. Nat. Struct. Mol. Biol. 19, 1316–1323 (2012).

48. Sloan, K. E., Knox, A. A., Wells, G. R., Schneider, C. & Watkins, N. J. Interactions and activities of factors involved in the late stages of human 18S rRNA maturation. RNA Biol. 16, 196–210 (2019).

49. Maurice, F., Pérébaskine, N., Thore, S. & Fribourg, S. In vitro dimerization of human RIO2 kinase. RNA Biol. 16, 1633–1642 (2019).

50. Agudelo, D. et al. Marker-free coselection for CRISPR-driven genome editing in human cells. Nat. Methods 14, 615–620 (2017).

51. Schmidt, E. K., Clavarino, G., Ceppi, M. & Pierre, P. SUnSET, a nonradioactive method to monitor protein synthesis. Nat. Methods 6, 275–277 (2009).

52. Nieto, B. et al. Identification of distinct maturation steps involved in human 40S ribosomal subunit biosynthesis. Nat. Commun. 11, 156 (2020).

53. Fischer, U. et al. A non-canonical mechanism for Crm1-export cargo complex assembly. Elife 4, (2015).

54. Geerlings, T. H., Faber, A. W., Bister, M. D., Vos, J. C. & Raué, H. A. Rio2p, an evolutionarily conserved, low abundant protein kinase essential for processing of 20 S Pre-rRNA in Saccharomyces cerevisiae. J. Biol. Chem. 278, 22537–22545 (2003).

55. Knüppel, R. et al. Insights into the evolutionary conserved regulation of Rio ATPase activity. Nucleic Acids Res. 46, 1441–1456 (2018).

56. Ameismeier, M., Cheng, J., Berninghausen, O. & Beckmann, R. Visualizing late states of human 40S ribosomal subunit maturation. Nature 558, 249–253 (2018).

57. Wyler, E. et al. Tandem affinity purification combined with inducible shRNA expression as a tool to study the maturation of macromolecular assemblies. RNA 17, 189–200 (2011).

58. Schäfer, T. et al. Hrr25-dependent phosphorylation state regulates organization of the pre-40S subunit. Nature 441, 651–655 (2006).

59. Ghalei, H. et al. Hrr25/CK1δ-directed release of Ltv1 from pre-40S ribosomes is necessary for ribosome assembly and cell growth. J. Cell Biol. 208, 745–759 (2015).

60. Zemp, I. et al. CK1δ and CK1ε are components of human 40S subunit precursors required for cytoplasmic 40S maturation. J. Cell Sci. 127, 1242–1253 (2014).

61. Mills, E. W. & Green, R. Ribosomopathies: There’s strength in numbers. Science 358, (2017).

62. Kressler, D., Hurt, E. & Bassler, J. Driving ribosome assembly. Biochim. Biophys. Acta 1803, 673–683 (2010).

63. Strunk, B. S. & Karbstein, K. Powering through ribosome assembly. RNA 15, 2083–2104 (2009).

64. Baumas, K. et al. Human RioK3 is a novel component of cytoplasmic pre-40S pre-ribosomal particles. RNA Biol. 9, 162–174 (2012).

65. Widmann, B. et al. The kinase activity of human Rio1 is required for final steps of cytoplasmic maturation of 40S subunits. Mol. Biol. Cell 23, 22–35 (2012).

66. O’Donohue, M.-F., Choesmel, V., Faubladier, M., Fichant, G. & Gleizes, P.-E. Functional dichotomy of ribosomal proteins during the synthesis of mammalian 40S ribosomal subunits. J. Cell Biol. 190, 853–866 (2010).

67. Zhao, Y., Bjorbaek, C. & Moller, D. E. Regulation and interaction of pp90(rsk) isoforms with mitogen-activated protein kinases. J. Biol. Chem. 271, 29773–29779 (1996).

